# SPICE: A Robust Computational Framework for Identifying Copy Number Variations in Spatial Transcriptomics

**DOI:** 10.64898/2026.06.30.735508

**Authors:** Kalins Banerjee, Robert C. Langefeld, Evan T. Keller, Xiang Zhou

## Abstract

Copy number variation (CNV), which alters the number of genomic segments, is a major driver of intratumor heterogeneity, characterized by spatially organized and genetically distinct cell populations. Recent advances in spatially resolved transcriptomic (SRT) technologies, which profile gene expression across thousands of spatially indexed tissue locations, offer a powerful opportunity to reconstruct the CNV architecture and dissect the spatial organization of cancer subclones. Here, we introduce SPICE (**sp**atial **i**nference of **C**NV **e**vents), a probabilistic method for identifying somatic CNVs and allele-specific copy number (ASCN) profiles from SRT data. A key feature of SPICE is its ability to integrate multiple complementary information available in SRT data, including gene expression, spatial coordinates, and heterozygous SNPs inferred from transcriptomic reads, to substantially enhance the accuracy and power of CNV detection. Using datasets generated across different SRT platforms, we first assess the reliability of SNPs derived from SRT data to ensure robust downstream inference. We then demonstrate that SPICE effectively integrates these modalities to deliver accurate and spatially coherent reconstruction of CNV landscapes and subclonal architecture, while maintaining excellent control of false discoveries. Together, SPICE provides a robust and effective solution for dissecting genomic heterogeneity in SRT studies of cancer.

## 1 Introduction

Intratumor heterogeneity is a fundamental feature of cancer, and is largely driven by genomic alterations including CNVs which are duplications and deletions of genomic regions [1]. CNVs promote cancer development and progression through amplification of oncogenes and deletion or inactivation of tumor suppressor genes [2]. In cancer tissues, CNVs can either be inherited or arise somatically [3], and become spatially segregated [2,4], giving rise to genetically distinct subclones that often differ in proliferation capacity, metastatic potential, and resistance to chemotherapy [2]. Characterizing the CNV architecture of cancer cells and mapping the spatial distribution of the associated subclones are therefore essential for understanding tumor evolution and heterogeneity, microenvironmental interactions, as well as treatment response.

Single-cell DNA sequencing (sc-DNAseq) remains the gold-standard for resolving per-cell CNV profiles [5], while emerging spatial DNA sequencing technologies provide direct insights into the spatial organization of cancer subclones [6]. However, sc-DNAseq is costly and often suffers from limited coverage, and spatial DNAseq technologies are still under active development. In contrast, a wide variety of RNAseq data, including bulk, single cell, and spatial, have become abundant and relatively straightforward to produce, which have motivated computational strategies to infer CNVs from these data. Such computational approaches rely on the assumption that genes located on amplified or deleted genomic segments often exhibit concordant shifts in expression levels relative to normal diploid genomic regions [4]. Building on this assumption, several methods [7–16] have been developed to infer CNVs from bulk or sc-RNAseq data.

Despite this progress, two major challenges remain. First, most existing RNAseq-based CNV inference methods were developed for bulk or sc-RNAseq studies, and are not suitable for the emerging SRT technologies. SRT platforms now enable transcriptomic profiling on tissues with spatial localization information, offering an unprecedented opportunity to map CNVs within their native spatial context. Although sc-RNAseq-based CNV inference methods are frequently used on SRT data [17–20], these methods do not account for the spatial information that is critical for characterizing heterogeneity and organization of cancer subclones [21].

Additionally, SRT data are, in general, more sparse as compared to sc-RNAseq data, especially with regard to SNP coverage, and thus contain relatively limited information for CNV inference [9,21,22]. Second, using only gene expression for CNV inference can be suboptimal. Specifically, gene expression is influenced by multiple regulatory mechanisms beyond CNV (e.g., regulation by transcription factors, DNA methylation, and chromatin modification), introducing substantial noise into CNV inference [13]. While matched DNA sequencing data are often limited, RNAseq data alone contain allelic information from SNPs, which can inform associated changes in allele-specific copy number (ASCN) i.e., number of copies of the haplotype chromosomes. Consequently, incorporating RNAseq-derived SNP information can, in principle, provide complementary evidence to gene expression shifts, and can substantially improve the resolution and robustness of CNV inference.

Although several recent tools [21–25] have been proposed for inferring CNVs from SRT data, they remain limited: many rely solely on gene expression, are designed for a specific SRT platform, or require matched DNA sequencing data. For example, SlideCNA [21] is designed specifically for Slide-seq gene expression data [26], whereas CalicoST [22] is designed primarily for Visium. Recently developed Clonalscope [25] does not explicitly model spatial dependencies among neighboring locations and requires allelic information from matched DNA sequencing data which most SRT experiments do not collect. As SRT technologies rapidly advance, there is an urgent need for a unified and platform-agnostic framework that can jointly model gene expression, spatial structure, and allelic information to robustly identify CNVs across diverse SRT platforms.

To address the above critical limitations, we present SPICE, a probabilistic framework, to identify somatic CNVs originating in cancer cells and associated ASCNs from SRT dataset. SPICE is compatible with SRT platforms of different resolutions which include multi-cellular resolution platforms such as Visium and near-single-cell resolution platforms such as Slide-seqV2 [27]. SPICE first identifies spatial domains within the malignant region, providing a spatially aware foundation for subclone detection. Then, it employs a Gaussian mixture model (GMM) for gene expression, a Binomial mixture model for allele counts at heterozygous SNPs, and finally, a data-adaptive joint likelihood framework integrating results from the two modalities. By bridging the gap between transcriptomic profiles and underlying DNA alterations in a spatially aware framework, SPICE offers an efficient and robust tool for dissecting the complex CNV architecture of SRT datasets from cancer tissues.

## 2 Results

### 2.1 Overview

We propose SPICE, a probabilistic method that integrates gene expression and allele counts from SNPs to identify genomic regions with CNVs in cancer tissues using SRT datasets. The proposed method is applicable for both spot level (multi-cellular resolution) and single-cell level (cellular resolution) SRT datasets. SPICE is designed to estimate CNVs by contrasting data from locations within the cancer region, often referred to as the malignant region, against those in the control region, referred to as the benign region. It takes four inputs: the standard UMI count matrix (gene x location), allele-count matrices for a set of heterozygous SNPs capturing read counts on reference and alternative alleles (SNP x location), coordinates of the tissue locations, and annotations that categorizes each location as either benign or malignant.

In the first step, we identify potential subclones within the malignant region using location-wise annotations and spatial domains derived with SpatialPCA, a spatially aware dimension reduction tool [28] (Figure 1, upper panel). SpatialPCA extracts a low dimensional representation (spatial principal components or spatial PCs) of the SRT dataset while preserving biological signal and spatial correlation structure. The spatial domains are then identified by clustering the tissue locations based on their spatial PCs. As a result, the tissue is segmented into multiple spatially organized and functionally distinct structures or microenvironments each of which is characterized by a distinct transcriptomic profile. We define a subclone as the collection of malignant locations belonging to a particular spatial domain. Thus, the malignant region of the tissue either can be represented by a single spatial domain (a single subclone) or can get divided into multiple subclones displaying transcriptomic heterogeneity. In the subsequent steps, we focus on identifying CNVs from each subclone using gene expression and allele counts from SNPs (Figure 1, lower panel).

**Figure 1.**
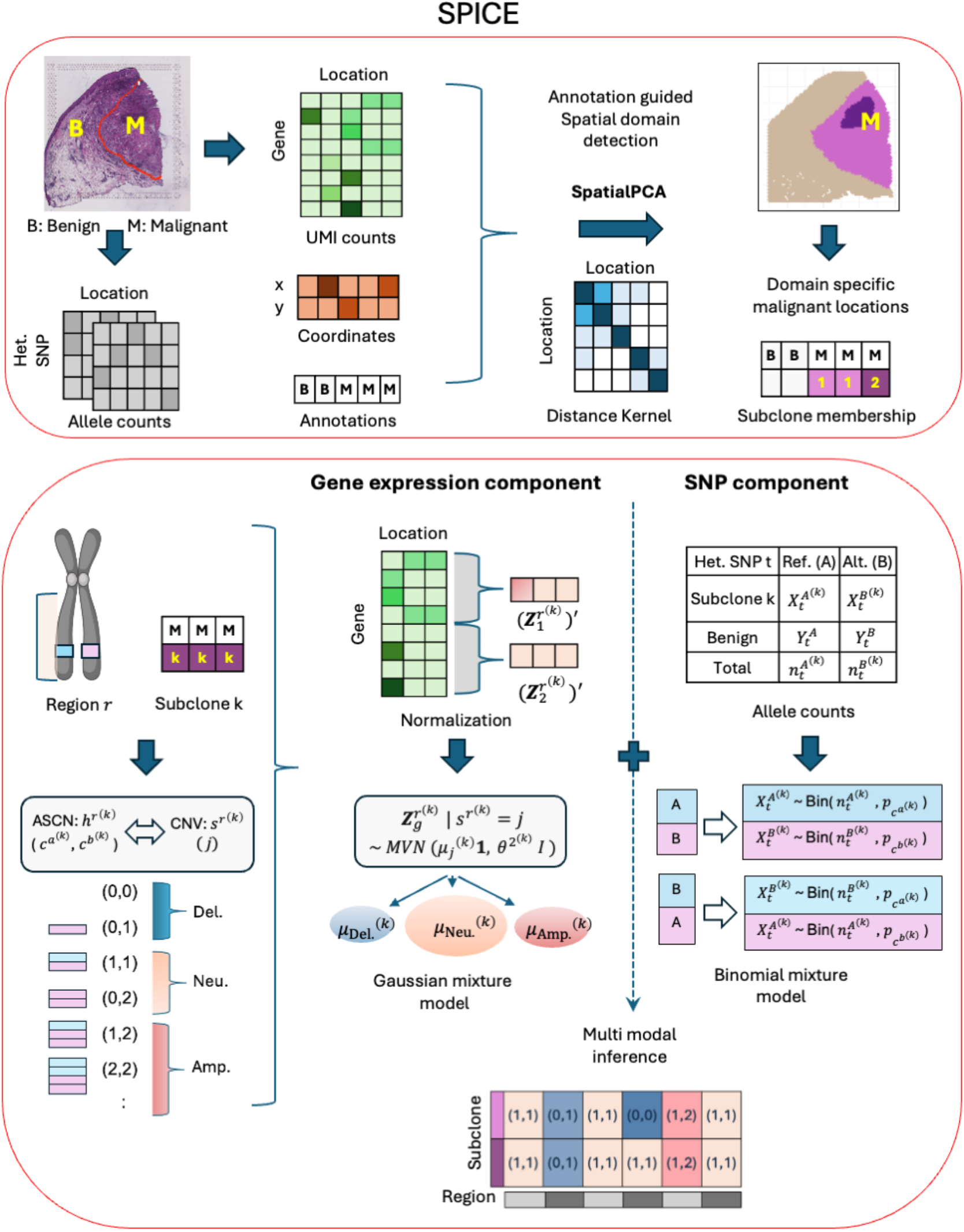
Schematic overview of SPICE. **Upper panel**: SPICE **(sp**atial **i**nference of **C**NV **e**vents) is a probabilistic method that identifies genomic regions harboring somatic CNVs from SRT data of cancer tissues. SPICE takes four inputs: UMI count matrix, allele-count matrices for a set of heterozygous SNPs, coordinates of the tissue locations, and annotations categorizing each location as either benign (B) or malignant (M). In the first step, SPICE identifies potential subclones within the malignant region based on location-wise annotations and spatial domains derived using SpatialPCA, a spatially aware dimension reduction tool which builds a kernel matrix to explicitly model the spatial correlation structure. **Lower panel**: For the gene expression component, a Gaussian mixture model is used for each subclone to represent the likelihood of normalized gene expression data under different CNV states. For the SNP component, a mixture of Binomial distributions is used for each subclone to identify CNV clusters based on ASCNs. Finally, the two modalities are combined in a joint likelihood to obtain posterior probabilities for CNV assignments.

For the gene expression component of our framework, we first normalize the gene expression data and perform baseline adjustment (section 4.1) so that any remaining relatively large-scale variations, observed in the malignant section of the tissue can be, on an average, attributed to the underlying CNVs. Then, a GMM is used for each subclone to represent the likelihood of the data under different CNV states (section 4.2). For the SNP component of our framework, we first obtain allele counts for a set of common, germline, bi-allelic, and heterozygous SNPs (section 4.3). We then aggregate the allele counts across each subclone as well as across the benign locations, to account for sparsity in coverage. Next, a mixture of Binomial distributions is used to model the allele counts and identify CNV clusters for each subclone based on ASCNs (section 4.4). Here, we follow the intuition that, for any allele, the proportion of reads mapped to a subclone among the reads mapped either to the benign region or to the subclone is, on an average, an increasing function of the number of copies of the underlying halpotype chromosomes in the subclone [29]. Finally, we combine results from the two modalities in a joint likelihood and obtain posterior probabilities for CNV assignments along the genome (section 4.5).

### 2.2 Assessment of SNPs derived from SRT datasets

For our research, we considered a human colon cancer dataset [6], generated using Slide-seqV2 [27], and two human cutaneous squamous cell carcinoma (cSCC) datasets [30], generated using the Visium platform (Additional details). Specifically, we considered cSCC datasets of patient 4 and patient 6 from the original study [30], following earlier publications [24,31]. We first investigated the extent to which germline SNPs could be recovered from SRT datasets. For ground truth, we obtained germline SNPs from matched single-cell whole-genome sequencing (scWGS) data associated with the colon cancer study and from matched whole-exome sequencing (WES) data associated with the cSCC study (section 4.6). We used bcftools [32] for scWGS data and Strelka2 [33] for WES data. The resulting variants which were also present in a panel of common SNPs (minor allele frequency > 0.05) from phase 3 of the 1000 Genomes Project were designated as germline (Additional details, Supplementary Figure 3) [34]. We referred this panel as ‘known SNPs’ for convenience. We considered Strelka2, bcftools, and cellsnp-lite [35], three widely used variant-callers in sc-RNAseq studies [9,10,36], to obtain germline SNPs from SRT samples (section 4.6). To the best of our knowledge, the existing studies had used cellsnp-lite to obtain SNPs from SRT datasets [9,22].

We first focused on the homozygous SNPs for evaluation (Figure 2 C & G). For this purpose, we compared SNPs detected as homozygous in a SRT sample with the homozygous SNPs identified from the corresponding benchmark dataset. We found that the average recall was only about 5% (range: 3.6-9.3%), whereas the precision levels varied greatly among the variant callers (range: 20.1-83.4%). For every dataset, cellsnp-lite achieved the best recall and the best F1 score, while Strelka2 had the best precision. Cellsnp-lite had relatively poor precision (avg. P= 21%), and the results from cellsnp-lite were quite similar whether all the locations or only the benign locations were used as input (section 4.6).

**Figure 2.**
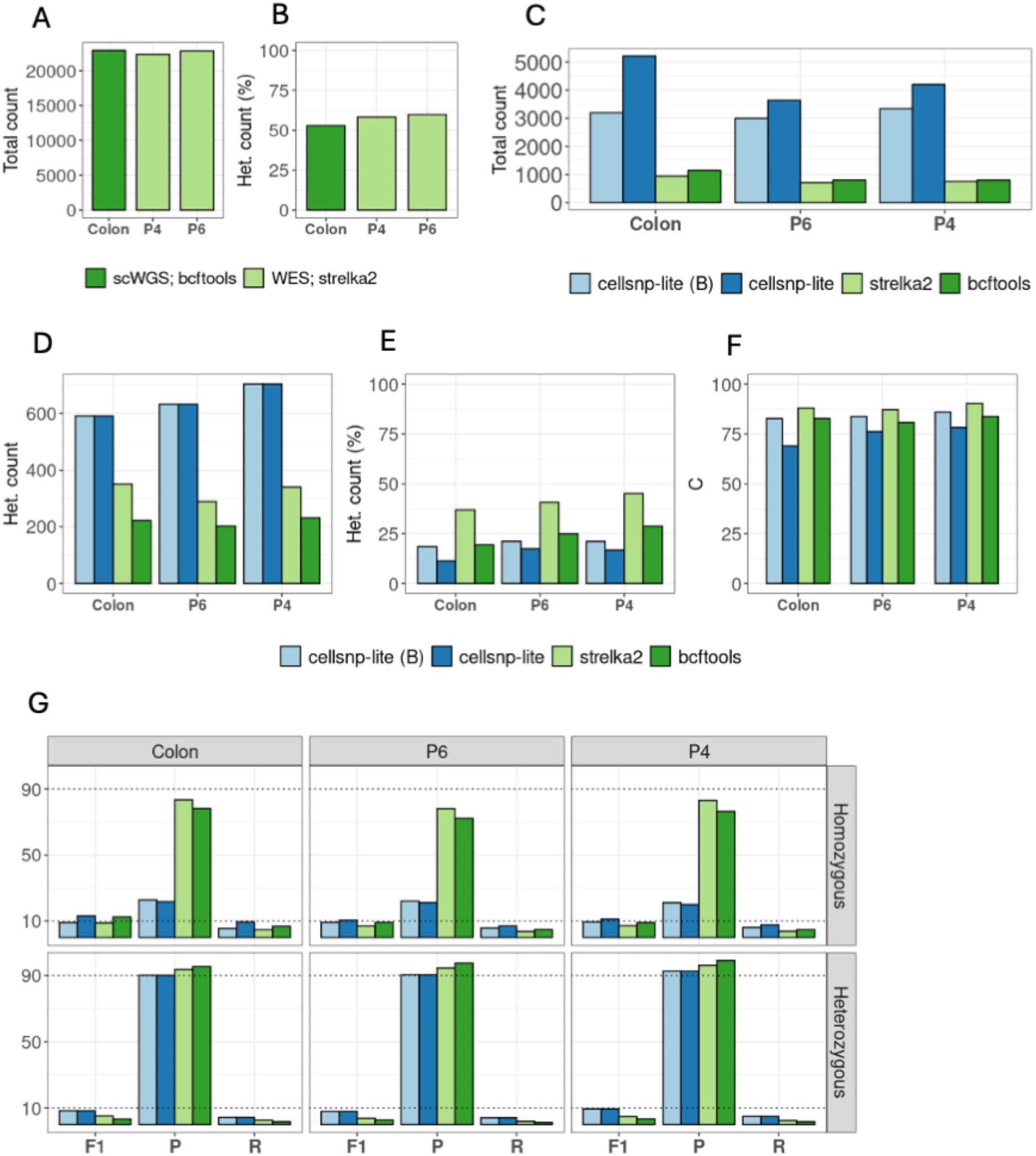
Assessment of the quality of germline, autosomal, and bi-allelic SNPs called from SRT samples. **A-B**. Details on SNPs serving as ground truth. Total SNP count (A) and percentage of heterozygous SNPs (B). Strelka2 was used for the WES samples and bcftools was used for the scWGS sample. **C-F**. Details on SNPs obtained from SRT samples. Total SNP count (C), heterozygous SNP count (D), percentage of heterozygous SNPs (E), and concordance rates (F). In ‘cellsnp-lite (B)’ the input bam file contained only the benign locations. **G**. F1 score, precision and recall (in percentages). Horizontally dotted lines mark 10 and 90. Legends are same as those for D-F. Colon: colon cancer study, P4: patient 4 from cSCC study, P6: patient 6 from cSCC study.

Next, we evaluated the quality of heterozygous SNPs (Figure 2 C-E, G). All three variant-callers achieved more than 90% precision (range: 90.2-99.1%), while the highest attained recall was only 5% (range: 1.4-5%). Clearly, a small proportion of germline heterozygous SNPs were recovered, likely due to the low RNA sequencing coverage from SRT platforms, but the identified SNPs were extremely rich in true positives. Among the three variant callers, bcftools had the best precision, while cellsnp-lite had the highest recall and the highest F1 score. Furthermore, bcftools identified the least number of heterozygous SNPs (avg. count=219), while cellsnp-lite had the highest number of calls (avg. count=642). Cellsnp-lite produced identical results regardless of whether all the locations or only the benign locations were used. The relatively high precision laid the foundation for incorporating heterozygous SNPs towards detecting CNVs from SRT datasets. In addition, given that cellsnp-lite achieved the best F1 scores and readily provided location-level allele counts (SNP x location matrix), we used it with all the tissue locations in our workflow.

While precision, recall, and F1 score were used to assess location-level matches within each genotype class, the concordance rate was used to examine all the location-level matches with respect to allele-level match and genotype-level match (Figure 2 F). Strelka2 had the best concordance rate (avg. C=88.5%). For cellsnp-lite, the approach involving only benign locations provided better results as compared to the approach involving all the locations (avg. C=84% vs 74.5%). We conducted two additional quality assessment studies. First, we examined the sensitivity of the findings with respect to the usage of the panel of known SNPs (Supplementary Figure 4). We found that the known SNPs acted as a filter in the final selection step, significantly improving the signal (true detection) to noise ratio. Specifically, we noticed a 44% drop in precision, on an average, for heterozygous SNPs without considering the panel. Second, we evaluated the quality of phased heterozygous SNPs, that were derived with a population based phasing technique [37], and had been used for CNV detection from sc-RNAseq or SRT datasets [9,10,22] (Supplementary Figure 3). The results revealed that the average precision was only 47.3%, while average recall was 3.5%. The details of the two studies can be found in the Supplementary Notes.

### 2.3 Application to human colon cancer Slide-seqV2 dataset

We benchmarked SPICE against established tools, designed for detecting CNVs from scRNA-seq and SRT data, including Numbat [9], CopyKAT [7], inferCNV [8], and CalicoST [22]. We further assessed the efficacy of the proposed joint inference framework by evaluating SPICE against its components: SPICE_GE (GMM applied to gene expression; section 4.2) and SPICE_SNP (Binomial mixture model applied to SNPs; section 4.4). For ground truth, we considered CNVs that were detected in matched DNAseq datasets (scWGS/WES), and were reported either in the original study or in published literature (section 4.7).

We assessed SPICE’s performance, starting with the colon cancer Slide-seqV2 dataset. SPICE achieved the best F1 score (Figure 3 E; F1=72.73%, P=66.67%, R=80%), followed by the F1 scores of Numbat and CopyKAT which were at least 60% lower (F1=28.57%, 20.09% respectively). We could not apply CalicoST as the Slide-seqV2 dataset was too sparse for the pipeline (https://github.com/raphael-group/CalicoST/issues/18). SPICE detected the least number of false CNVs by achieving the best precision. Numbat had the second-best precision (P=50%), while precisions of other methods were less than 30%. In terms of retrieving true CNVs, inferCNV had the best performance (R=90%). But it had the poorest precision (P=11.54%) which led the F1 score to be almost 72% lower than the best. We further compared the accurate CNV calls of SPICE with those of Numbat and CopyKAT, the two best competing methods. It turned out that all the true detections of Numbat and CopyKAT (amplifications on 8q, 20p and 20q) were identified by SPICE. Additionally, SPICE had several unique recoveries (deletions on 15q, 18p and 18q; amplifications on 1q and 7p).

**Figure 3.**
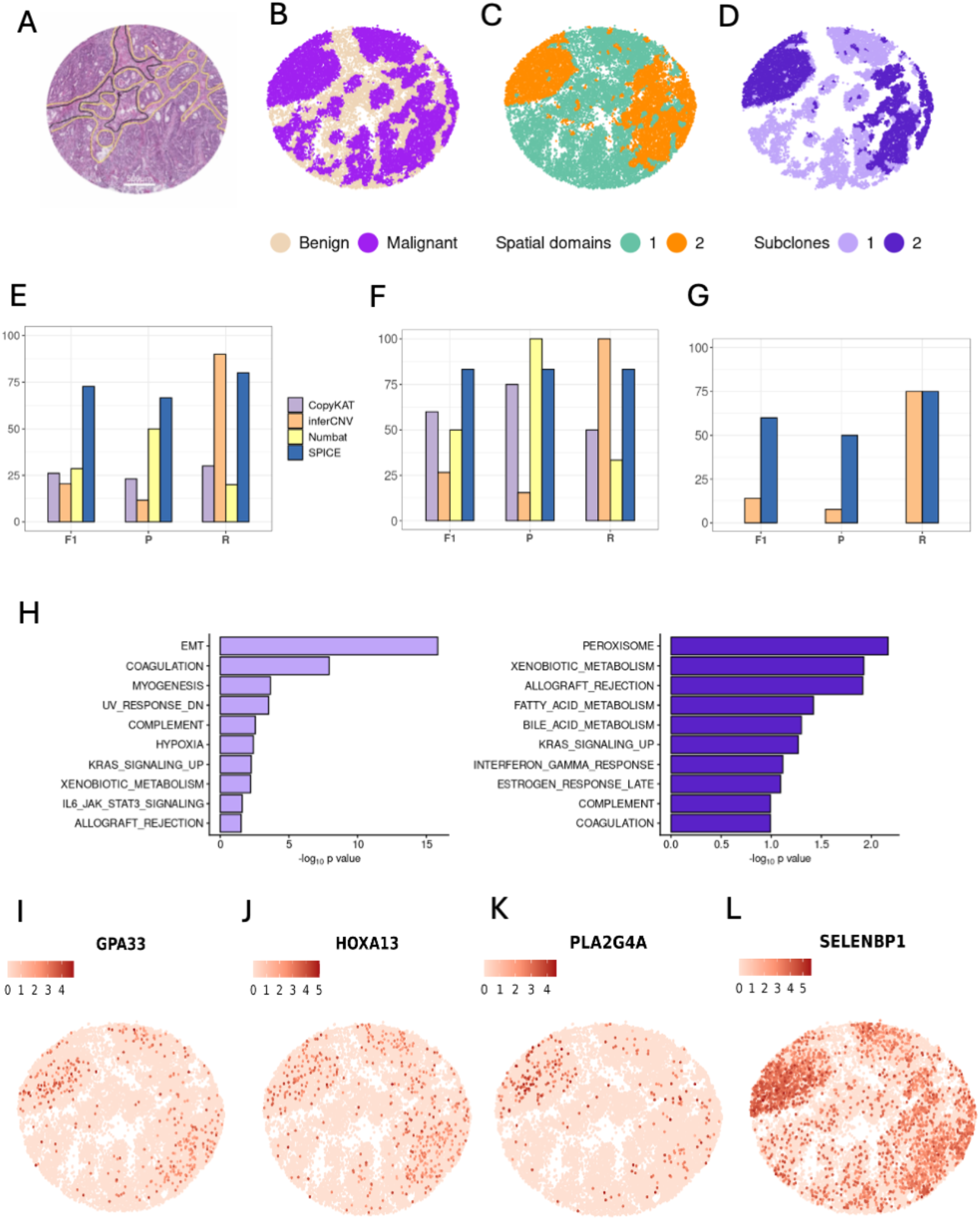
Analysis of colon cancer Slide-seqV2 dataset. **A**. H&E image of the tissue obtained from [49]. **B-D**. Scatter plot showing annotated locations (B) and spatial domains with number of clusters set as 2 (C). SPICE partitioned the malignant locations into two subclones (D). **E**. F1 score, precision and recall (in percentages) comparing true CNVs (deletions and amplifications) and identified CNVs. **F-G**. F1 score, precision and recall (in percentages) comparing true amplifications vs. identified amplifications (F) and true deletions vs. identified deletions (G). The legends are same as those in E. **H**. Top 10 enriched gene sets from GSEA in subclones identified by SPICE (left: subclone 1; right: subclone 2). The Hallmark gene sets linked to specific biological pathways were used. EMT: epithelial–mesenchymal transition. **I-L**. Examples of genes characterizing subclone 2. Differential expression analysis was performed between the two subclones, and DE genes were mapped to the CNV harboring regions successfully traced by SPICE. The plotted genes were upregulated in subclone 2 and had well-established roles in colorectal cancer biology. The normalized expressions were plotted (layer=‘data’) using Seurat::FeaturePlot().

Next, we considered the two CNV classes (deletion and amplification) separately to examine the robustness of the performances. SPICE was able to achieve the best F1 scores in both classes (Figure 3 F-G). It retrieved around 83% amplifications with 83% precision. The second best F1 score in this class belonged to CopyKAT (F1=60%). Interestingly, Numbat did not make any false amplification calls, while inferCNV retrieved all the amplifications. But Numbat’s recall and inferCNV’s precision were relatively poor. On the other hand, only SPICE and inferCNV were able to identify true deletions. While both achieved 75% recall, SPICE outperformed inferCNV with a substantially higher precision (P= 50% vs. 7.69%) that ultimately provided it with the best F1 score within the deletion class.

We next evaluated the efficacy of SPICE’s joint modelling framework by examining the performances of its components. SPICE_GE had a much higher recall than SPICE_SNP (R= 80% vs. 30%), while the latter achieved perfect precision (P=100%). SPICE efficiently integrated the CNV-signals from the two modalities (Supplementary Figure 5 A-B). First, it preserved all the true detections of its components. Second, SPICE rectified multiple false detections (e.g., chromosomes 1, 4, 5, 7q, 10p, 12q, and 19) of SPICE_GE without making additional false detections. Consequently, SPICE was able to outperform its components.

SPICE can further characterize the diversity within the malignant region by identifying subclones along with the corresponding clonal (shared) and subclonal (subclone-specific) CNVs. In this dataset, SPICE identified two spatially segregated subclones (Figure 3 D), of almost equal size (47% and 53% of locations respectively). SPICE revealed the subclonal nature of multiple amplifications. To illustrate, subclone 1 was characterized by amplification on chromosome 20p, while subclone 2 was characterized by the co-occurrence of amplifications on chromosomes 1q and 7p (Supplementary Figure 5 A). Similar subclonal amplifications from two subclones were also reported in the original study (Fig.3e of [6]). SPICE also accurately identified multiple CNVs shared by the subclones (deletions on 15q, 18p, and 18q; amplification on 8q). It missed the subclonal nature of a single CNV (20q amplification) among its accurate calls. We further analyzed the matched scWGS dataset and found that SPICE accurately estimated ASCNs for several of its true CNV calls (e.g., one copy amplification on 1q, 7p, 20p, and 20q; detailed in Supplementary Notes).

As a next step, we performed differential expression analysis between the two subclones to examine changes in gene expression within the inferred subclonal CNVs. We identified multiple upregulated genes, having well-established roles in colorectal cancer (CRC) biology, from the subclonally amplified regions (Figure 3 I-L, Supplementary Figure 5 C-H). Examples from subclone 2 included upregulated expression of HOXA13, SCIN, SELENBP1, LGR6, PLA2G4A, LEFTY1, and GPA33. HOXA13 (7p), a prognostic biomarker for CRC, reportedly facilitates metastasis by transactivating ATP-citrate lyase and insulin-like growth factor 1 receptor [38]. SCIN (7p) is known for its elevated expression in CRC patients with synchronous liver metastasis and a poor prognosis [1]. LGR6 (1q) plays a tumor-promoting role in CRC development, and its elevated expression has been associated with tumor differentiation [39]. On the other hand, the aggressiveness of CRC reportedly increases with decreased SELENBP1 (1q) expression [40]. PLA2G4A (1q) is also noteworthy given its sidedness-dependent prognostic association-higher expression correlating with improved overall survival in left-sided CRC but poorer prognosis in right-sided CRC [41]. GPA33 (1q) has been recognized for its heterogeneous expression in CRC with antigen loss at the infiltrative tumor edge [42]. Finally, LEFTY1 (1q) is a robust biomarker for tumor-initiating cells in colonic tissues [43].

Conversely, among the DE genes upregulated in subclone 1, RALGAPA2 and NOP56 were located on chromosome 20p, a segment exhibiting amplification in this subclone. Higher expression of NOP56 has been associated with adverse outcome in overall survival of patients with advanced colon cancer [44], while RALGAPA2 has been found upregulated in sporadic CRC and in normal colon as compared to colitis-associated cancer [45]. To gain further insights of the identified subclones, we performed gene set enrichment analysis (GSEA) of the differentially expressed (DE) genes using the Hallmark gene sets linked to specific biological pathways [46] (Figure 3 H). We found that subclone 1 was enriched in multiple pathways including epithelial mesenchymal transition, coagulation, myogenesis, complement system, hypoxia, ultraviolet response (down), and KRAS activation (up). Meanwhile, subclone 2 was enriched in pathways of peroxisome and fatty acid metabolism. Notably, genes supporting xenobiotic metabolism and allograft rejection were upregulated in both subclones.

### 2.4 Application to human squamous cell carcinoma Visium datasets

In this section, we present results from the two Visium datasets, starting with patient 6. SPICE demonstrated outperformance by achieving the best F1 score (Figure 4 E; F1=48%, P=85.7%, R=33.3%). Among the competing methods, inferCNV, Numbat, and CopyKAT had similar F1 scores (range= 35.71-37.68%). SPICE achieved the best precision which was almost 54% higher than the next-best precision of Numbat (P=55.56%). SPICE had the second-best recall while inferCNV had the best recall at the cost of poorest precision (P=25.49%; R=72.22%).

We next compared the methods by considering amplifications and deletions separately (Figure 4 F-G). SPICE identified amplifications with the best F1 score and the best precision (F1=66.7%, P=83.3%), while Numbat was the best competitor in these two criteria. Although inferCNV achieved the best recall, its precision was the poorest. On the other hand, methods relying solely on gene expression, such as CopyKAT and inferCNV, outperformed within the deletion class. It turned out that methods incorporating the SNP modality were extremely conservative in identifying deletions. For instance, SPICE made a single correct deletion call (13q) without inferring any false deletions (F1=20%).

**Figure 4.**
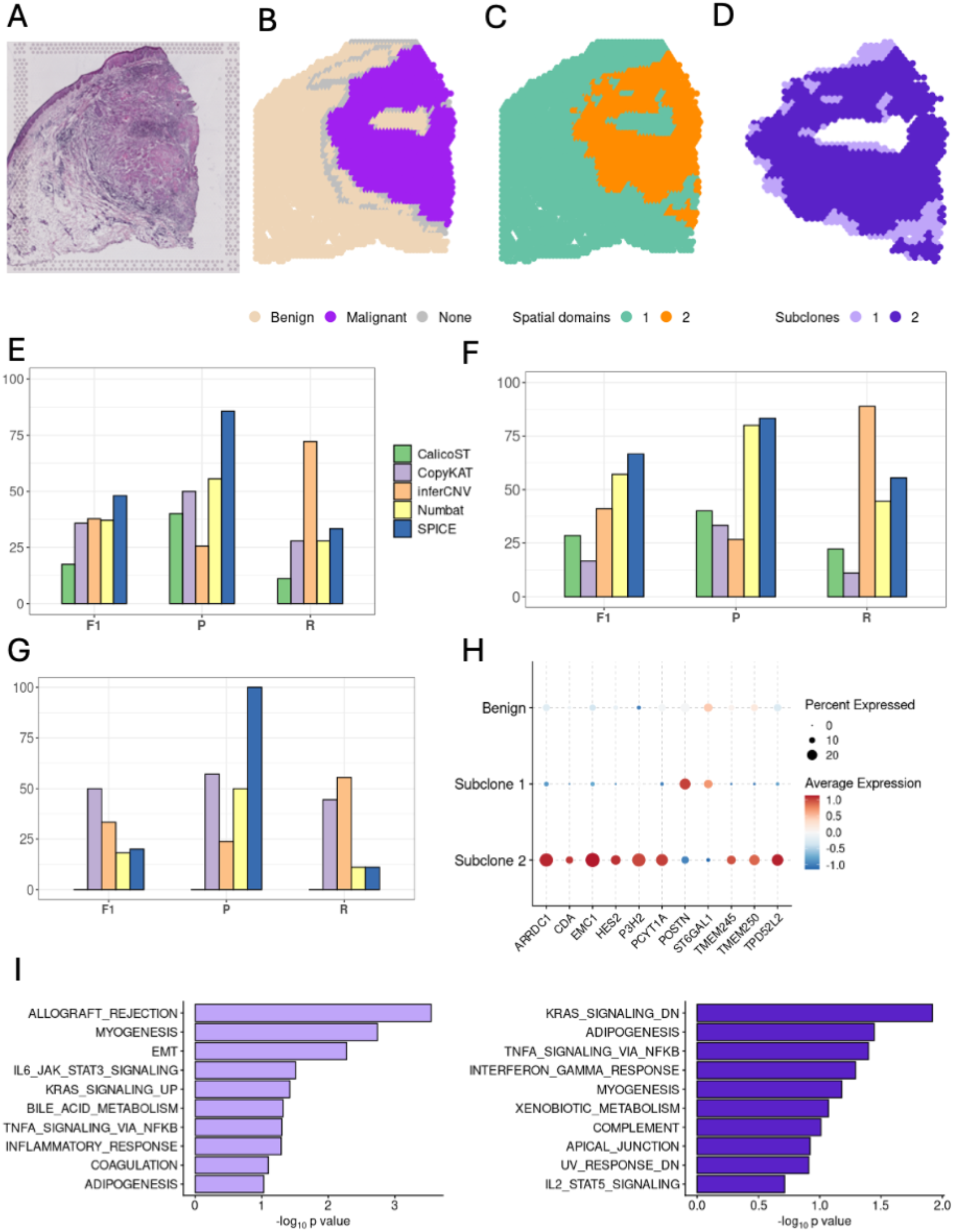
Analysis of patient 6 Visium dataset. **A**. H&E image of the tissue. **B-D**. Scatter plot showing annotated locations (B) and spatial domains with number of clusters set as 2 (C). SPICE partitioned the malignant locations into two subclones (D). **E**. F1 score, precision and recall (in percentages) comparing true CNVs (deletions and amplifications) and identified CNVs. **F-G**. F1 score, precision and recall (in percentages) comparing true amplifications vs. identified amplifications (F) and true deletions vs. identified deletions (G). The legends are same as those in E. **H**. Dot plot showing examples of genes characterizing the two subclones. Differential expression analysis was performed, and DE genes were mapped to the CNV harboring regions successfully traced by SPICE (CDA, HES2, & EMC1 are on 1p. P3H2, ST6GAL1, & PCYT1A are on 3q. TMEM250, ARRDC1, & TMEM245 are on 9q. TPD52L2 is on 20q). POSTN (13q) showed reduced expression in subclone 2 where 13q had undergone deletion. The size of the dot encodes the percentage of locations within a class. **I**. Top 10 enriched gene sets from GSEA in the two subclones (left: subclone 1; right: subclone 2). The Hallmark gene sets linked to specific biological pathways were used. EMT: epithelial–mesenchymal transition.

We further investigated the relatively weak recall of SPICE in identifying deletions by examining the performances of its components (Supplementary Figure 6 E-F). SPICE_GE outperformed SPICE_SNP by a great margin (F1= 77.78% vs. 28.58%), whereas SPICE_SNP, demonstrating perfect precision, identified three amplifications (1p, 3q, and 20q). It turned out that, SPICE was unable to preserve multiple true deletions (3p, 4q, 10q, 21q) along with some true amplifications (8q, 14q, 17q) identified by SPICE_GE. Still, it benefited from the perfect precision of SPICE_SNP as it overturned many false detections from SPICE_GE (deletions on 5p; amplifications on 11p and 16p). Overall, SPICE demonstrated robustness by smoothing out the contrasting performances of its components and produced superior results.

We next investigated whether the CNVs were spread across the malignant region or prevalent within some subclones. SPICE partitioned the malignant region into two subclones (Figure 4 C-D), where subclone 1 had only 226 locations and existed primarily along the boundary of the malignant region. Interestingly, SPICE did not detect any CNVs in this subclone. Thus, all the detections of SPICE belonged to subclone 2. We further validated the corresponding ASCNs by analyzing the matched WES dataset. SPICE was accurate in predicting one copy amplification with most of its accurate amplification calls (Supplementary Notes).

Next, we performed DE analysis between the two subclones to compare gene expression patterns with a particular focus on the CNV harboring regions successfully traced by SPICE (Figure 4 H, Supplementary Figure 7). Among the DE genes mapped to regions harboring amplification in subclone 2, PCYT1A (3q; upregulated), has been highlighted as a candidate gene in squamous cell carcinoma (SCC) biology, including reports of tumor-suppressive effects in lung SCC and head and neck SCC, where higher PCYT1A expression correlates with improved survival [46,47]. Another example involved upregulation of ARRDC1(9q). Published evidence suggests context-dependent roles for ARRDC1, including reported downregulation with tumor-suppressive activity in clear cell renal cell carcinoma [48] and upregulation in hepatocellular carcinoma [49]. Conversely, POSTN (13q) showed reduced expression in subclone 2, consistent with the observed 13q deletion. In cSCC, POSTN has been reported to be highly expressed in both malignant and stromal compartments [47]. Besides, elevated expression of POSTN is associated with poor prognosis in many cancers including cSCC [47,48]. GSEA analysis with the Hallmark gene sets linked to specific biological pathways further found that subclone 1 was enriched in pathways related to epithelial mesenchymal transition, allograft rejection, myogenesis, KRAS activation (up), and IL-6/JAK/STAT3 signaling (Figure 4 I). On the other hand, subclone 2 was enriched in pathways of KRAS activation (down), adipogenesis, and TNFA signaling via NF-κB.

Next, we focus on findings from the Visium dataset of patient 4 where SPICE had the best performance as it accurately estimated all chromosome arms to have ASCN of (1,1) (Supplementary Figure 8). Here, the SNP modality had a substantial impact in reducing the number of false detections to zero as SPICE_SNP had no false detections while SPICE_GE had 10 false calls. Besides, the malignant locations belonged to a single spatial domain. Among the competing methods, CopyKAT had a single false call (amplification on 18q), while Numbat and CalicoST had two false calls each (Numbat: deletion on 1q and 5q; CalicoST: copy number neutral loss of heterozygosity on 1q and deletion on 5q). Surprisingly, inferCNV identified 47 CNV-harboring segments, spanning the entire autosome. Notice that, inferCNV had the poorest precision also in the other real data applications.

Finally, it is worth highlighting that SPICE’s integration framework is flexible in assigning more weightage to a specific modality. Accordingly, preference to a particular modality based on prior knowledge can be easily incorporated. For instance, the two Visium datasets we analyzed had, on an average, twice the number genes on each chromosome arm than the colon cancer Slide-seqV2 dataset (avg. gene count= 363 vs. 740; Supplementary Figure 1 A-B). Besides, the average total UMI count and the average number of genes expressed at each location were also higher for the Visium samples (Supplementary Figure 1 E-G). On the other hand, the average total read count per heterozygous SNP was lower on each arm for the Visium dataset of patient 6, reflecting lower confidence on the SNP modality (Supplementary Figure 2 B). We utilized this observation by assigning 25% additional weightage to SPICE’s gene expression component (*w*=0.75; section 4.5) for the Visium datasets. As a result, there was a substantial boost in SPICE’s ability to retrieve true CNVs from the patient 6 dataset at a minor compromise in precision (F1= 60%, P=75%, R= 50%). Specifically, SPICE was able to identify two additional deletions (3p and 5q) and one additional amplification (8q) in subclone 2, further enhancing its superiority (Supplementary Figures 6G & 10). At the same time, the superior result of zero CNV calls from the patient 4 dataset remained unaffected (Supplementary Figure 11). We additionally conducted an extensive sensitivity analysis with respect to the weighting strategy and demonstrated SPICE’s robust performance across all the real datasets considered (Supplementary Notes, Supplementary Figures 9-11).

## 3 Discussion

We introduced SPICE, a probabilistic framework designed to identify somatic CNVs from SRT sample of cancer tissues. A key strength of SPICE is its compatibility and robust performance across different spatial resolutions. The efficacy of SPICE was elucidated through its application to real-world datasets from two commercialized SRT platforms: Visium Spatial Gene Expression by 10x Genomics, a platform having multi-cellular resolution, and Slide-seqV2, a near-single-cell resolution platform, commercialized as Curio Seeker. A significant contribution of our research also included the systematic assessment of germline SNPs derived from SRT datasets. To our knowledge, this represented the first such evaluation, revealing a substantially high signal-to-noise ratio in heterozygous SNPs. Even though we encountered poor recalls, the high precision levels provided a strong foundation for using heterozygous SNPs to identify CNVs and infer the associated ASCNs. SPICE’s flexible joint likelihood framework efficiently integrated gene expression and allele counts to provide superior performance while ensuring excellent protection against false detections. The GMM, applied to gene expression, exhibited an edge in retrieving true CNVs, whereas the Binomial mixture model, designed for heterozygous SNPs, primarily served as a safeguard against false detections. SPICE efficiently modelled this complementary relationship to demonstrate robust performance.

Addressing the intrinsic challenges of SRT data, such as high sparsity or zero inflation, was a primary focus in our development. We observed significant sparsity in both gene expression (locations with zero total UMI; avg.= 97%, range=72-99%; Supplementary Figure 1) and in heterozygous SNP data (locations with zero total allele count; avg.= 99.5%, range=80-100%; Supplementary Figure 2). We accounted for sparsity in two ways. First, we focused on relatively large scale CNVs by considering chromosome arms as analysis unit. Second, we aggregated sparse allele counts within each subclone and within the benign region for the Binomial mixture model. Additionally, we emphasize the importance of expert-guided annotations in our framework. SPICE requires annotated benign and malignant locations as inputs. This requirement is often fulfilled by pathologists examining the H&E images associated with SRT experiments. Existing CNV detection pipelines can separate benign and malignant locations using different approaches that lack extensive benchmarking. Besides, a SRT location can be composed of multiple cells (e.g., 1-10 cells for Visium), and SRT data, collected along the boundary of the malignant region, are often composed of a mixture of malignant and benign cells. These locations can introduce errors into the analysis, providing unreliable biological conclusions. In fact, identifying malignant region from muti-cellular SRT data is a separate area of research [49]. Thus, we preferred annotations provided by a pathologist, often available along with the published dataset. Otherwise, marker gene expression, deconvolution tools, which have been comprehensively benchmarked [50], or external pipelines [49] might be used for annotation.

Methodologically, SPICE leverages SpatialPCA-derived spatial domains and annotated malignant locations to identify subclones. The spatial domains segment the tissue into multiple spatially organized and functionally distinct structures having unique transcriptomic profiles. When restricted within the malignant region, these distinct transcriptomic profiles of the spatial domains can reflect the underlying CNV profiles. Our approach of identifying subclones accounts for the spatial correlation structure of the tissue-a feature that is often ignored by existing pipelines. Since SpatialPCA has been recognized as a top-performing tool for spatial domain detection in terms of accuracy and robustness [51], its integration into our framework ensures that subclone identification is both accurate and biologically grounded. Besides, any efficient and robust spatial domain detection tools [51] can be easily incorporated into the proposed framework.

Despite the capabilities, SPICE has multiple areas for future expansion. Currently optimized for a single tissue slice of an SRT experiment, the framework could be extended to incorporate multiple slices or to modalities other than SRT. Considering the high sparsity of SRT datasets, integration with features extracted from the accompanying H&E images, or integration with matched sc-RNAseq datasets, have the potential to amplify the underlying CNV signals. Finally, SPICE is designed for SRT platforms compatible for variant calling. For instance, allele counts cannot be derived from imaging based or probe-based platforms such as Xenium, FFPE compatible Visium or Visium HD. Although the gene expression component of SPICE can be applied to these datasets, such an application would carry higher risk of false detections.

In conclusion, SPICE represents a robust, resolution independent, and spatially aware method for characterizing the CNV architecture of SRT datasets. We expect that the two modalities used in SPICE would complement each other in providing robust results across a wide range of datasets through the proposed joint inference framework.

## 4 Methods

### 4.1 Preprocessing of gene expression data

The UMI count matrix (gene x location) was subjected to multiple preprocessing steps prior to the analysis. Initially, cycle genes, HLA genes, non-autosomal genes, and mitochondria genes were left out, following standard practice [7]. Next, genes expressed in fewer than 5 locations, or with total UMI count less than 100, or that displayed extreme variability (more details below) were not considered. Finally, locations with total UMI count less than 100 were filtered out. In order to account for variation in sequencing depths, the UMI counts were then scaled so that the total UMI count of each location became equal to the median of the location-wise total UMIs. Subsequently, log transformation was applied using a pseudo count of 1.

A standard preprocessing step for identification of CNVs from RNA-seq data is to smooth out gene-level variations by considering a group of neighboring genes as unit of analysis instead of individual genes [23,52]. We followed a similar procedure to construct a matrix (bin x location) of normalized expression by computing successive averages over bins, comprised of 100 neighboring genes, along the autosomes. In order to create a baseline for each bin, we then computed the corresponding average normalized expression over the set of benign locations. Finally, the baseline expression of each bin was subtracted from its normalized expression at each malignant location, and the log transformation was inverted by exponentiation [8,23].

We noticed certain characteristic differences among the gene expression (UMI count) datasets generated using different platforms (e.g., Visium, Slide-seqV2). Such characteristics included proportion of genes expressed at each location, total UMI count from each location, and variation in gene expression across locations (Supplementary Figure 1). Accordingly, the preprocessing pipeline included certain steps that were specific to the data generating platform. In particular, we used the 95^th^ and 99^th^ percentiles of gene-wise variances for Visium and Slide-seqV2 data respectively as thresholds to identify genes with extreme variation.

### 4.2 Gaussian mixture model for gene expression data

Let there be *n* many malignant locations *n*^(*k*)^ of which belong to subclone *k*. For any bin *g* located on chromosome arm *r*, let 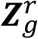 denote the resulting vector with elements representing baseline adjusted and normalized expressions of malignant locations. Henceforth, we refer the elements of 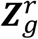 as normalized expressions for simplicity. Let 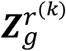 be a sub-vector of 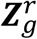 having elements representing locations from subclone *k*. Our objective from SRT gene expression data is to estimate the most likely CNV state of each chromosome arm in the cancer region. The CNV state of any region in the genome (e.g., a chromosome arm) is characterized by the underlying number of parental halpotype chromosomes, and the corresponding total chromosome count is defined as the total copy number of that region. There are three major CNV states, namely deletion, copy number neutral, and amplification, where deletion refers to a state having total copy number less than 2, copy number neutral refers to a state having 2 as total copy number, and any state with total copy number greater than 2 is considered as an amplification state. A diploid normal human genome possesses a single copy of each of the two parental haplotype chromosomes. Consequently, the total copy number of any region in a diploid normal human genome is 2. On the other hand, each of the two haplotype chromosomes underlying any CNV-affected region on a cancer genome either gets deleted or amplified (i.e., duplicated) by at least one copy. Furthermore, if a genomic region harbors a CNV during cancer development, the CNV can either be prevalent across the entire malignant region (i.e., clonal CNV) or be limited to only a subsection of the malignant region (i.e., subclonal CNV). As a result, a CNV affected cancer genome can be uniquely partitioned within each subclone into multiple genomic regions where each region can be assigned to one of the three major CNV states. Notice that, a copy number neutral state indicates that the underlying region either has not been affected by any CNVs or has undergone one copy deletion along with one copy amplification (i.e., copy number neutral loss of heterozygosity). Even though RNAseq gene expression data can be used to separate genomic regions having different total copy numbers, it cannot distinguish among the different configurations generating same total copy number (e.g., the two configurations generating copy number neutral state).

Considering the high sparsity level in typical SRT gene expression datasets, we limit our gene expression-based CNV detection to the three major states. In other words, we assume that the underlying signal might not be rich (or dense) enough to distinguish between two copy deletions and a single copy deletion, or between a single copy amplification and various multi-copy amplifications. Due to the same reason, we have focused on estimating relatively large scale CNVs at the chromosome arm level. Accordingly, let 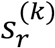 denote the true CNV state of chromosome arm *r* within subclone *k*. For ease of presentation, we denote the three CNV states by integers (1: deletion, 2: copy number neutral, 3: amplification). We model the conditional distribution of 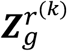 given the underlying CNV state 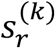 as a mixture of three multivariate Normal distributions while applying a hierarchical prior on 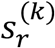. Specifically,

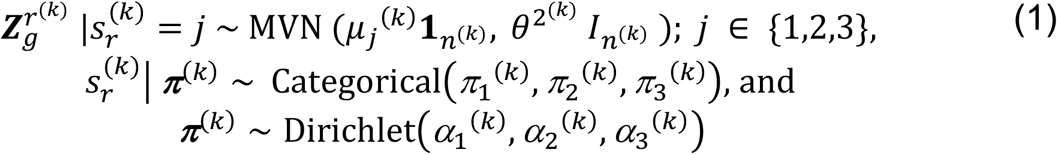

Here, 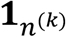 is a vector of *n*^(*k*)^ many ones, 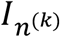 is an identity matrix of order 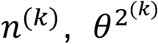 is the subclone specific variance component, and *μ*_*j*_ ^(*k*)^ is the subclone specific expected normalized expression of a bin harboring CNV state *j* . Since, the normalized expressions are expected to be directly proportional to the underlying total copy numbers on an average, we get *μ*_1_^(*k*)^ < *μ*_2_^(*k*)^ < *μ*_3_^(*k*)^. In practice, we have fixed 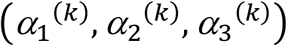 as (0.05, 0.99, 0.05), which assigns a high prior belief for a region to remain copy number neutral. This is consistent with the fact that most regions, in general, are not expected to have undergone any CNVs. Besides, as shown in results (sections 2.3-2.4), such a stringent assignment has provided SPICE with substantial protection against false detections. We have estimated the parameters 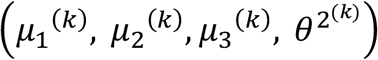 using the Expectation-Maximization (EM) algorithm. Further details on parameter estimation are provided in the Supplementary Notes. We denote the results obtained from this GMM as SPICE_GE.

### 4.3 Preprocessing of SNP data

We followed an existing approach [9] to obtain a set of germline, bi-allelic, autosomal, and heterozygous SNPs. Specifically, we used cellsnp-lite [35], a tool for genotyping bi-allelic SNPs, with a panel of known common SNPs (minor allele frequency > 0.05) from the 1000 Genomes Project (Additional details). As a result, we obtained a matrix (SNP x location) of alternative allele counts, representing the number of reads mapped to the alternative alleles, and another matrix of the same dimension having total counts i.e., total number of reads mapped to the reference and alternative alleles. Then, we applied a few quality control steps. First, we only considered SNPs with positive total count over the set of benign locations. Then, we eliminated SNPs with excessively high total counts using the 99^th^ percentile of the SNP-wise total counts as threshold. Locations with zero total count combining all SNPs were also eliminated. Finally, SNPs with proportion of alternative allele counts within the range of [0.10, 0.90] over the set of benign locations were identified as germline heterozygous [9].

### 4.4 Binomial mixture model for SNPs

Let the two alleles at each heterozygous SNP be denoted as *A* and *B*, and the two halpotype chromosomes be labelled as *a* and *b*. We define, 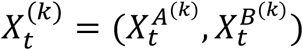, as allele counts at heterozygous SNP *t* from locations in subclone *k*. Additionally, let 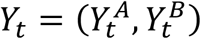, be allele counts at the same heterozygous SNP *t* from locations in the benign region. Next, for each allele, we define the total counts combining locations from subclone *k* and the benign region as 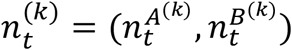, where 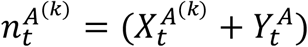 and 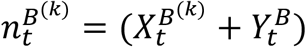. The phasing information i.e., whether a specific allele is located on chromosome *a* or chromosome *b*, remains usually unknown. Let *I*_*t*_ be a latent indicator that equals 1 if allele *A* is situated on chromosome *a* and equals 0 if allele *A* is situated on chromosome *b*. Since, Mendel’s law of equal segregation allows the allele to have an equal chance to be located on any one of the two haplotype chromosomes, we have the prior probability *P*[*I*_*t*_ = 1] = 0.5 [29]. Besides, the value of *I*_*t*_ remains same across all the cells.

Haplotype-specific copy number or allele-specific copy number (ASCN) of any genomic region is defined as the number of copies of the underlying haplotype chromosomes. A diploid normal human genome possesses a single copy of each of the two haplotype chromosomes. Consequently, the ASCN for any genomic region can be represented as (1,1). On the other hand, CNVs divide the cancer genome into several contiguous genomic regions that are characterized by the underlying ASCNs. For example, a single copy from both arm 1p and arm 2p might undergo deletion, while both the copies of arm 1q might remain intact. In this case, the ASCNs of the three contiguous arms (1p, 1q, and 2p) can be represented as (0,1), (1,1) and (0,1) respectively. As any somatic CNV can either be prevalent across the entire malignant region or be limited to only a subsection of the malignant region, the ASCNs of a CNV-affected cancer genome can be represented by piece-wise constants within each subclone.

Let 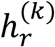 be a variable denoting the ASCN of genomic region *r* (e.g., a chromosome arm) within subclone *k*, and let 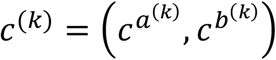 denote an observed value of 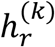, where 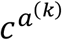 and 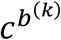 respectively represent the number of copies of chromosomes *a* and *b* underlying that region within subclone *k*. Additionally, for any CNV affected region, we denote the haplotype chromosome with lesser number of copies as chromosome *a*. Thus, *c*^(*k*)^ = (0,1) represents deletion of chromosome *a*, whereas *c*^(*k*)^ = (1,1) represents a region that has not undergone any CNVs. Now, for each heterozygous SNP *t* in region *r*, we use a pair of Binomial distributions to model the allele counts from subclone 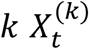, conditional on the combined allele counts 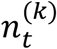, the phasing indicator *I*_*t*_, and the underlying ASCN 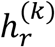. Specifically,

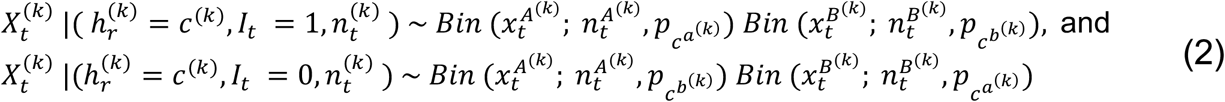

Here, the notation *Bin*(*x*; *n, p*) denotes a Binomial distribution with mean *np*, and *x* is an observation of the underlying random variable. As the copy number of a haplotype chromosome underlying any region of the cancer genome increases (or decreases), if we consider reads mapped to an allele from that region and reads mapped to the same allele from a matched normal diploid genome, the proportion of the former over the total is expected to increase (or decrease) on an average. In other words, the Binomial parameter 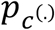 can be expected to behave like an increasing function of the corresponding ASCN *c*^(.)^. Besides, modelling the conditional distribution of the allele counts from malignant locations given the allele counts combining malignant and benign locations allows us to internally control for various experimental influences. Similar modelling approaches have been used earlier in the context of inferring ASCNs from whole genome sequencing data [29] and whole-exome sequencing data [53]. Since we model each subclone separately, we drop the superscript (*k*) henceforth for ease of notation. Then, the joint distribution of SNPs can be written as 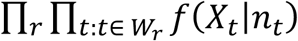 where

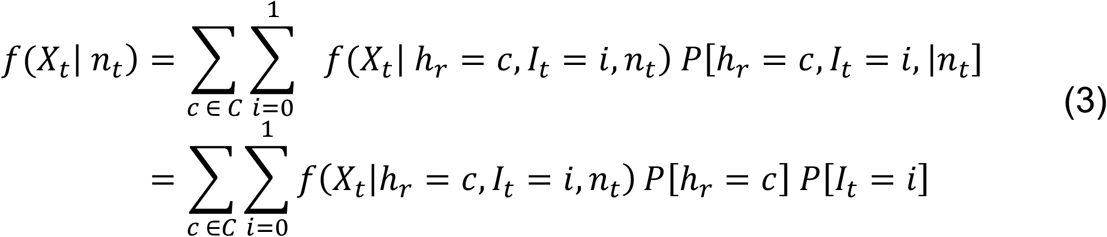

Here, *W*_*r*_ denotes the set of heterozygous SNPs in region *r, C* denotes the set of ASCNs, and the distribution *f*(*X*_*t*_|*h*_*r*_ = *c, I*_*t*_ = *i, n*_*t*_) has been specified in (2). Considering the ASCNs that are both likely to be inferred from RNA-seq experiments, and have been selected in existing literature, we have specified *C* as {(0,0), (0,1), (1,1), (0,2), (1,2), (1,3), (2,2)} [9]. Similar to a recent study [54], our modelling approach treats each chromosome arm independently while inferring the underlying ASCNs. Besides, we have considered the chromosome arms as genomic regions considering the sparse SNP coverage of typical SRT datasets. We have used an EM algorithm to estimate the Binomial parameters (*p*_0,_ *p*_1,_*p*_2,_*p*_3_), and the corresponding details are provided in the Supplementary Notes. We denote the results obtained from this Binomial mixture model as SPICE_SNP.

### 4.5 Joint inference from gene expression and SNPs

The GMM, described in section 4.2, can identify the most likely CNV state (deletion, copy number neutral, amplification). However, it cannot provide further information regarding the underlying ASCNs. In addition, it might be difficult to distinguish among the deletion and amplification levels that result in different total copy numbers (e.g., two copy deletions, one copy deletion, one copy amplification, two copy amplifications etc.) due to the relatively high sparsity in SRT gene expression datasets. On the other hand, the Binomial mixture model, described in section 4.4, can provide the most likely copy number configurations at the haplotype level. However, its performance may be compromised by the sparse SNP coverage in SRT datasets. Therefore, we propose SPICE, a likelihood-based framework, that efficiently integrates results from the two modalities, to provide accurate and robust results across a wide variety of SRT datasets. Our objective is to assign an ASCN *c*_0_ to each chromosome arm, such that, *c*_0_ maximizes the posterior assignment probability conditional on two data modalities. Mathematically,

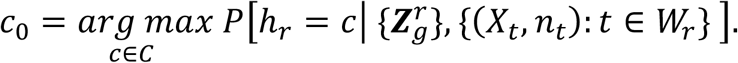

Now, if we assign a fractional weight of *w* ∈ (0,1) to the gene expression component, the objective function takes the following form.

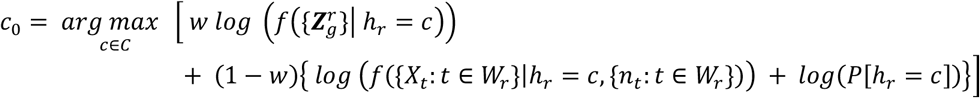

Derivation of the objective function along with the computational details are provided in Supplementary Notes. Unless stated otherwise, we set *w* = 0.50, assigning equal weightage to both modalities. Finally, CNV states are assigned to chromosome arms based on posterior probabilities. Specifically, we first obtain the most likely ASCN (*c*_0_), and then convert it to the corresponding CNV state. For instance, configurations such as (1,2), (2,2), and (1,3) are classified as amplifications, while (0,0) and (0,1) are classified as deletions.

### 4.6 Assessment of SRT SNPs

The motivation for considering SNPs to identify CNVs is well established. Even though it appears intuitive that incorporating SNPs into the CNV-analysis of SRT data would be beneficial, a critical first step involves assessing the quality of the SNPs. Although multiple studies have assessed the quality of SNPs obtained from sc-RNAseq [20,21], to the best of our knowledge, there exists no such study aimed at SRT data. In line with our objective, we investigated the extent to which germline SNPs could be recovered from SRT datasets. Besides, it was imperative to quantify the signal-to-noise ratio in the derived SNP-set.

For the colon cancer study, we treated germline SNPs obtained from the scWGS data as ground truth to assess the quality of those obtained from the matched Slide-seqV2 data. We used bcftools [32] for variant calling from scWGS, and restricted the calls to lie within the exonic regions of the genome (Additional details). The variants which were also present in a panel of common SNPs (minor allele frequency > 0.05) from phase 3 of the 1000 Genomes Project were designated as germline (Additional details, Supplementary Figure 3) [34]. We have referred this panel of SNPs as ‘known SNPs’ in this article for convenience. For the cSCC study, we analyzed samples from patient 4 and patient 6, following earlier publications [24,31]. The study also included WES data from the tumor and the adjacent normal skin. We obtained germline variants from the matched WES of normal skin using Strelka2 (Additional details) [33,55]. Then, variants, present also in the panel of known SNPs, were considered as ground truth while assessing the quality of germline SNPs obtained from the corresponding Visium samples (Supplementary Figure 3).

In order to obtain germline SNPs from SRT samples, we selected widely used variant-callers from sc-RNAseq studies [9,10,36]. Specifically, we considered Strelka2, bcftools, and cellsnp-lite [35] (Additional details, (Supplementary Figure 3). It is worth noting that, to the best of our knowledge, cellsnp-lite is the only variant caller that had been used to obtain SNPs from SRT datasets without any quality-assessment [9,22]. Since, cellsnp-lite had the convenience of extracting location-level allele counts from a single bam file, we examined two approaches involving cellsnp-lite to obtain germline SNPs. In the first approach, we used the original SRT bam file as input. Then, SNPs with proportion of alternative allele counts within the range of [0.10, 0.90] over the set of benign locations were identified as germline heterozygous [9]. In the second approach, we used a subsetted bam file, retaining only the benign locations, as input. The criterion for identifying heterozygous SNPs remained unchanged. Unless mentioned otherwise, results from cellsnp-lite would refer to the first approach involving all the tissue locations. As opposed to cellsnp-lite, Strelka2 and bcftools did not readily provide the location-level allele counts from a single bam file. Thus, we used the bam file, having only benign locations, as input. We identified heterozygous variants for these two pipelines based on their default genotyping. The variant calls from SRT data were restricted within the exonic regions of the genome for fair assessment (Additional details). Lastly, we selected variants, present also in the panel of known SNPs, as candidates. Although only heterozygous SNPs were used in SPICE, we investigated the quality of germline homozygous SNPs for the purpose of completeness. We characterized a SNP by its location in the genome, and used precision (P), recall (R), F1 score, and concordance rate (C) as evaluation criteria. A true detection referred to a SNP that had been detected both in a SRT dataset and in the corresponding benchmark dataset. Precision measured the percentage of true detections among the SNPs called from a SRT sample, recall measured the percentage of true detections among the SNPs called from a benchmark dataset, and F1 score summarized precision and recall through the harmonic mean. Finally, the concordance rate measured the percentage of true detections where the alleles (reference and alternative) and the genotype (heterozygous/ homozygous) got matched.

### 4.7 Evaluation of SPICE

The colon cancer study [6] used matched slide-DNA-seq data to identify CNVs which were further validated based on matched scWGS data. The study reported four deletions (15p, 15q, 18p, and 18q) and six amplifications (1q, 7p, 7q, 8q, 20p, and 20q) where some amplifications (1q, 7p, 7q, 20p, and 20q) were subclonal in nature [6]. We used these reported CNVs as ground truth. For the cSCC study [30], a recent publication [31] analyzed the WES datasets from patient 4 and patient 6 for CNVs, and we used the corresponding results as ground truth for our research. Interestingly, no CNVs were detected in patient 4. On the other hand, several arm level CNVs were identified in patient 6. Specifically, nine deletions (3p, 4q, 5q, 10p, 10q, 13p, 13q, 21p, and 21q) and nine amplifications (1p, 3q, 8q, 9q, 11q, 14q, 17q, 20p, and 20q) were reported (Fig. S1 in [31]). We used precision (P), recall (R), and F1 score as evaluation criteria for datasets with known CNVs (colon cancer and patient 6). Precision measured the percentage of true CNVs among the detected CNVs and recall measured the percentage of true CNVs that were detected, among all the true CNVs. For patient 4, we considered the number of false detections as evaluation criterion. For each competing method, we used the malignant locations for CNV detection and counted all CNVs detected on each arm (Additional details).

## 5 Additional details

### Application of SpatialPCA

We used SpatialPCA (version 1.3.0), as a part of the SPICE pipeline, to identify subclones within the malignant region of the tissue. The UMI count matrix and the coordinates were used as inputs to SpatialPCA. Following the tutorials and platform-specific guidelines, we applied the Louvain algorithm for the Slide-seqV2 sample, and the Walktrap algorithm for the Visium samples [28].

For the Slide-seqV2 sample, we considered findings from the original study [6] and set the desired number of clusters as 2. This resulted in two subclones as the malignant region was spread across two spatial domains. For the Visium samples, we examined two choices (2 and 3) as the desired number of clusters. In both cases, the distribution of spatial domains over the malignant region remained same, and as we increased the desired number of clusters from 2 to 3, the new cluster was contained within the benign region (Supplementary Figures 6 & 8). As a result, number of subclones remained unchanged. For the Visium dataset of patient 6, the malignant region was spread across two spatial domains, resulting in two subclones. On the other hand, all the malignant locations from the Visium dataset of patient 4 belonged to a single spatial domain.

### Colon cancer Slide-seqV2 dataset

The colon cancer Slide-seqV2 dataset [6] had 16,270 genes and 18,288 locations where 12,736 locations were annotated as tumor and 5,552 locations were annotated as stroma. We obtained the annotations from the authors of a recent publication [49]. We used the locations from the stroma region as controls (benign locations).

### cSCC Visium datasets

The patient 6 Visium dataset from the cSCC study [30] had 33,538 genes and 3,650 locations where 1,361 locations were annotated as tumor and 2,002 locations were annotated as non-tumor. The annotations were obtained from a recent study [49]. We treated the non-tumor locations as controls. The patient 4 visium dataset from the cSCC study [30] had 33,538 genes and 744 locations. We used the annotations shown in Supp. Fig. 18A of [56] and Fig. S3 of [31] to identify 155 malignant locations and 400 benign locations. We selected the first replicate for each of the two samples following existing publications [49,56].

### Colon cancer scWGS dataset

The colon cancer study [6] performed scWGS, in addition to Slide-seqV2, on the same sample. Briefly, nuclei were extracted from a 5 mm x 12 mm x 100 um colon cancer section, and protocols of 10X Chromium Single Cell DNA were followed for library generation and sequencing [6]. We analysed the bam file: ‘human_colon_cancer_3_scwgs_210402_02’ which was shared by the authors of the original study [6]. For obtaining germline SNPs, that were used as gold standard, exons were specified using the Agilent BED file (SureSelect Human All Exon V6 r2; hg19).

### cSCC WES datasets

The cSCC study included patient-matched whole-exome sequencing (WES) data from the tumor and the adjacent normal skin samples. In short, the Agilent V6 whole-exome capture kit (60 Mbp) was used, and the resulting libraries were sequenced on BGISEQ-500 using paired-end 100bp reads to achieve approximately 100x coverage for each tumor sample and 50x coverage for each normal skin sample [30]. The publicly available fastq files were downloaded from the European Nucleotide Archive (ENA) and were subjected to TrimGalore (version 0.6.7) for quality control [57]. The resulting fastq files were mapped to hg38 using the Burrows-Wheeler Aligner (BWA-MEM; version 0.7.17), and the duplicate reads were removed using samtools markdup (version 1.19.2). We used the same reference genome (hg38) that was used in Space Ranger for mapping the Visium samples. The variant callers were restricted to the exonic regions of the genome as per established guideline [57]. We specified the exonic regions through the Agilent target BED (SureSelect Human All Exon V6 r2; hg38) file.

### Panel of known SNPs

Following standard practice [9,10,22], we used a list of common SNPs (minor allele frequency > 0.05) from phase 3 of the 1000 Genomes Project [58] for identifying germline SNPs from SRT data. The same list was also used to select the germline SNPs, used as ground truth, while assessing the quality of SRT SNPs. This list was downloaded from cellsnp-lite’s website [35]. We used the hg38 based list for the cSCC datasets, and the hg19 based list for the colon cancer datasets. This selection was made in accordance with the reference genomes used to generate the corresponding bam files.

### cellsnp-lite

cellsnp-lite (version 1.2.3) was used with minMAF=0 and minCOUNT= 2 to derive germline SNPs from SRT data [9]. We also used a list of common SNPs (minor allele frequency > 0.05) from phase 3 of the 1000 Genomes Project (-R option). We further retained SNPs located within the exonic regions for the quality assessment study.

### Strelka2

We used the --exome option with WES bam files and the –rna option with SRT bam files (Strelka2 version 2.9.10). Additionally, the –callRegions option was used in both cases for restricting the variants to the target regions (BED file) of WES. The resulting vcf files were annotated using bcftools annotate and a list of common SNPs (minor allele frequency > 0.05) from phase 3 of the 1000 Genomes Project. Finally, we restricted our analysis to SNPs that passed the default filters (FILTER=PASS). We used the genotype labels (GT column in vcf) to identify heterozygous SNPs.

### bcftools

For SRT, we used bcftools mpileup (version 1.19) with options: -Q 30 -A -x -Ou -I -R [36] whereas in case of scWGS, we used bcftools mpileup with --ignore-RG -R [34]. In both cases, the -R option was used so that the variants could be restricted to the exonic regions, specified through BED files. As before, the resulting vcf files were annotated based on a list of common SNPs (minor allele frequency > 0.05) from phase 3 of the 1000 Genomes Project using bcftools annotate.

### CopyKAT

The benign spots were specified through the ‘norm.cell.names’ option (CopyKAT version 1.1.0). Since, CopyKAT does not explicitly call copy number events, we followed an established approach to infer the CNV state of each chromosome arm [9]. First, we computed average copy number intensities of each arm over the predicted aneuploid spots. Then, we used thresholds of -0.03 and 0.03 to identify deletions and amplifications respectively.

### inferCNV

The benign spots were specified using the ‘ref_group_names’ option within CreateInfercnvObject(). For CNV calling we selected denoise=T, cutoff=0.1, cluster_by_groups=TRUE, analysis_mode=subclusters, HMM_report_by=subcluster, and HMM_type=i6 (version 1.26.0) [9].

### Numbat

We followed the tutorials and set the max_entropy as 0.8 (version 1.5.1). We used the benign locations to create the reference with aggregate_counts(). We first obtained the clone assignments of the locations (‘clone_opt’ column in clone_post_2.tsv). Then we mapped the segments with CNVs to chromosome arms (‘seg’ column in joint_post_2.tsv). Finally, for each clone, we recorded the CNVs detected in each arm.

### CalicoST

We applied CalicoST (version 1.0.0) on the two Visium datasets. We followed the tutorial without the optional steps. We first mapped each genomic region (cnv_seglevel.tsv) to chromosome arms. Then, for each clone we recorded the CNVs identified on each arm. CalicoST was designed primarily for Visium samples. The colon cancer Slide-seqV2 dataset was too sparse for the pipeline. Additionally, it could not use location-wise annotations (https://github.com/raphael-group/CalicoST/issues/18).

### Differential expression analysis and gene set enrichment analysis

For differential expression analysis, we conducted Wilcoxon rank sum test by using FindMarkers() from Seurat (v5). We declared a gene to be an upregulated gene if its adjusted p value was less than 0.05 and its log fold change was positive. Similarly, we declared a gene to be a downregulated gene if its adjusted p value was less than 0.05 and its log fold change was negative.

We used fgsea (version 1.30.0) [59] with Hallmark gene sets linked to specific biological pathways [46] for gene set enrichment analysis (GSEA). Specifically, we used genes with positive log fold change as input to fgseaMultilevel().

## Data Availability

Publicly available Visium and WES data from the cSCC study [30] were obtained from Gene Expression Omnibus (GEO): GSE144240. The Slide-seqV2 and scWGS bam files from the colon cancer study [6] were shared by the authors of the original study [6]. The UMI count matrix, coordinates and location-wise annotations associated with the Slide-seqV2 dataset were shared by the authors of a recent publication [49].

## Code availability

The SPICE software code is publicly available at https://github.com/kzb193/SPICE. The source code is released under the GNU General Public License version 3.

## Acknowledgements

This study was supported by the National Institutes of Health (NIH) grants R01GM126553, R01HG011883, R01HG009124, and R01GM144960.

## Author contributions

X.Z. conceived the idea and provided funding support. K.B. and X.Z. designed the experiments. K.B. developed the method and analyzed real data. K.B. and R.C.L. implemented the software. K.B., R.C.L., X.Z., and E.T.K. wrote the manuscript. All authors read and approved the final manuscript.

