## Supplementary Material for "SPICE: A Robust Computational Framework for Identifying Copy Number Variations in Spatial Transcriptomics"

### 3. Supplementary Notes

#### 3.1 Parameter estimation in GMM for gene expression data

This section provides an outline of the steps involved in fitting the GMM described in the main text. Since, we fit the GMM for each subclone separately, we drop the superscript $(k)$ henceforth for ease of notation. First, the complete data likelihood can be written as,

$$\prod_r f\left(\{\boldsymbol{Z}_g^r\}, s_r\right) = \prod_r \prod_{j \in \{1,2,3\}} f\left(\{\boldsymbol{Z}_g^r\}, s_r = j\right)^{1(s_r = j)}$$

$$= \prod_r \prod_j \left( f\left(\{\boldsymbol{Z}_g^r\} \mid s_r = j\right) P[s_r = j] \right)^{1(s_r=j)}$$

$$= \prod_r \prod_j \left( \{\prod_g f\left(\boldsymbol{Z}_g^r \mid s_r = j\right)\} \; P[s_r = j] \right)^{1(s_r=j)}$$

Here, $\{ \boldsymbol{Z}_g^r\}$ denotes the collection of bins within region $r$. Now, the complete data log-likelihood takes the following form.

$$log\ (\prod_r f(\{\boldsymbol{Z}_g^r\}, s_r))$$

$$= \sum_r \sum_{j=1}^{3} 1(s_r = j)\ (\sum_g log\ (f(\boldsymbol{Z}_g^r\ | s_r = j))\ +\ log\ (P[s_r = j])\ )$$

Notice that,

$$P(s_r = j) = (\int f(s_r = j|\boldsymbol{\pi}) f(\boldsymbol{\pi})\, d\boldsymbol{\pi}\ ) = \frac{\alpha_j}{\sum_l \alpha_l}$$

As described in the main text, we have fixed $(\alpha_1, \alpha_2, \alpha_3)$ as (0.05, 0.99, 0.05), assigning a high prior belief for a region to remain copy number neutral. Next, given a set of initial estimates for $(\mu_1, \mu_2, \mu_3, \theta^2)$, the E-step can be implemented as,

$$1(\widehat{s_r = j}) = P[s_r = j|\{\ \boldsymbol{Z}_g^r\}]$$

$$= \frac{P(s_r = j)(\prod_g f(\boldsymbol{Z}_g^r|\ s_r = j))}{\sum_{j_0}\ P(s_r = j_0)\ (\prod_g f(\boldsymbol{Z}_g^r|\ s_r =\ j_0))}$$

We have used the k-means algorithm for initial estimates of $(\mu_1, \mu_2, \mu_3)$, and we have set the initial estimate of $\theta^2$ as 1. Finally, at the M-step, $1(\widehat{s_r = j})$ is plugged into the complete log-likelihood which is then maximized.

#### 3.2 Parameter estimation in Binomial mixture model for SNPs

This section provides an outline of the steps involved in fitting the Binomial mixture model described in the main text. As the copy number of a haplotype chromosome underlying any region of the cancer genome increases (or decreases), if we consider reads mapped to an allele from that region and reads mapped to the same allele from a matched normal diploid genome, the proportion of the former over the total is expected to increase (or decrease) on an average. In other words, the Binomial parameter $p_{c^{(.)}}$ (main text; equation (2)) can be expected to behave like an increasing function of the corresponding ASCN $c^{(.)}$. Accordingly, we have initialized the Binomial parameters based on the relationship $p_{c^{(.)}} = \frac{c^{(.)}}{c^{(.)} + \alpha}$, where $\alpha$ is the ratio between number of reads mapped to the set of malignant locations and the set of benign locations. A proof of this relationship can be found in existing publications [1,2].

First, the complete data log-likelihood can be written as,

$$log(\prod_{r} \prod_{t:t \in W_r} f(X_t, h_r, I_t | n_t))$$

$$= \sum_{r} \sum_{t:t \in W_r} log\ (\prod_{c \in C} \prod_{i=0}^{1} (f(X_t | h_r = c, I_t = i, n_t) P[h_r = c]\ \ P[I_t\ = i])^{1(h_r = c, I_t = i)}\ )$$

$$= \sum_{r} \sum_{t: t \in W_r} \sum_{c \in C} \sum_{i=0}^{1} 1(h_r = c, I_t = i) \; \left[ log\left(f(X_t | h_r = c, I_t = i, n_t)\right) + log(P[h_r = c]) + log(P[I_t = i]) \right]$$

Then, the E-step can be computed as outlined below.

$$\widehat{1(h_r = c, I_t = i)} = P[h_r = c, I_t = i | X_t, n_t]$$

$$= \frac{f(X_t, h_r = c, I_t = i | n_t)}{f(X_t | n_t)}$$

$$= \frac{f(X_t | h_r = c, I_t = i, n_t) \; P[h_r = c, I_t = i | n_t]}{\Sigma_{c_0} \Sigma_{i_0} f(X_t | h_r = c_0, I_t = i_0, n_t) \; P[h_r = c_0, I_t = i_0 | n_t]}$$

$$= \frac{f(X_t | h_r = c, I_t = i, n_t) P[h_r = c] P[I_t = i]}{\Sigma_{c_0} \Sigma_{i_0} f(X_t | h_r = c_0, I_t = i_0, n_t) P[h_r = c_0] P[I_t = i_0]}$$

Finally, at the M-step, $\widehat{1(h_r = c, I_t = i)}$ is plugged into the complete log-likelihood which is then maximized with respect to $(p_{0,}\, p_{1,} p_{2,} p_3)$.

Next, we specify the prior probabilities that are kept fixed. Notice that, the probability that region $r$ remains copy number neutral is given as, $P[h_r = (1,1)] + P[h_r = (0,2)]$. Similarly, the probability of region $r$ undergoing deletion can be computed as, $P[h_r = (0,0)] + P[h_r = (0,1)]$, and the same for amplification can be computed as,

$P[h_r = (1,2)] + P[h_r = (1,3)] + P[h_r = (2,2)]$. Accordingly, we get the following relationships between the prior probabilities of the GMM (Note 3.1) and the Binomial mixture model.

$$P(s_r = 1) = (P[h_r = (0,0)] + P[h_r = (0,1)]) = 0.05,$$

$$P(s_r = 2) = (P[h_r = (1,1)] + P[h_r = (0,2)]) = 0.99,$$

$$P(s_r = 3) = (P[h_r = (1,2)] + P[h_r = (1,3)] + P[h_r = (2,2)]) = 0.05$$

Next, we have used fractional weights to connect prior probabilities of the ASCNs with prior probability of the corresponding CNV state. For example, the prior probabilities for the ASCNs corresponding to deletion are specified through the fractional weight $\zeta_{01}$ as,

$$P[h_r = (0,1)] = \zeta_{01} P(s_r = 1),$$

$$P[h_r = (0,0)] = (1 - \zeta_{01)} P(s_r = 1)$$

Similarly, for amplification, we use the fractional weight $\zeta_{12}$ as,

$$P[h_r = (1,2)] = \zeta_{12} P(s_r = 3);$$

$$P[h_r = (1,3)] = P[h_r = (2,2)] = \frac{1 - \zeta_{12}}{2} P(s_r = 3)$$

Finally, for CNV neutral states having total copy number as 2, we use fractional weight $\zeta_{02}$ as,

$P[h_r = (0,2)] = \zeta_{02} \ P(s_r = 2)$

$P[h_r = (1,1)] = (1 - \zeta_{02}) \ P(s_r = 2)$

We have set $\zeta_{01} = 0.60$, $\zeta_{12} = 0.60$, and $\zeta_{02} = 0.0001$ as the default. We have assigned a high prior belief for a region to remain copy number neutral, and such a stringent assignment has provided SPICE with substantial protection against false detections. We have further conducted a sensitivity analysis with respect to the choices for the fractional weights, and demonstrated SPICE's robust performance (Note 3.8).

#### 3.3 Computational details for joint inference in SPICE

Here, we describe computational details of SPICE's likelihood-based joint inference framework that combines the results from both data modalities. Our objective is to assign an ASCN $c_0$ to each genomic region ($r$) such that, $c_0$ maximizes the posterior assignment probability conditional on two data modalities. Mathematically,

$$c_0 = \underset{c \in C}{arg\,max} \ P\left[h_r = c \middle| \{\boldsymbol{Z}_g^r\}, \ \{(X_t, n_t): t \in W_r\} \right]$$

Now, the quantity on the right can be simplified as shown below.

$$\underset{c \in C}{arg\,max} \ P\left[h_r = c \middle| \{\boldsymbol{Z}_g^r\}, \{(X_t, n_t): t \in W_r\}\right]$$

$$= \underset{c \in C}{arg\,max} \;\; f\big( \{\boldsymbol{Z}_g^r\},\ \{X_t\!: t \in W_r\} \big|\ h_r = c,\ \{n_t\!: t \in W_r\}\big)\ P[h_r = c]$$

$$= \underset{c \in C}{arg\,max} \;\; f\big(\{\boldsymbol{Z}_g^r\}\big|\ h_r = c\big)\ f(\{X_t\!: t \in W_r\}|\ h_r = c,\ \{n_t\!: t \in W_r\})\ P[h_r = c]$$

Thus, the objective function can be equivalently written as,

$$c_0 = \underset{c \in C}{arg\,max} \;\; \big[log\ (f\big(\{\boldsymbol{Z}_g^r\}\big|\ h_r = c\big))$$

$$+\ log\ \big(f(\{X_t\!: t \in W_r\}|h_r = c, \{n_t\!: t \in W_r\})\big)\ +\ log(P[h_r = c])\big]$$

If we assign a fractional weight of $w \in (0,1)$ to the gene expression component, the objective function takes the following form.

$$c_0 = \underset{c \in C}{arg\,max} \;\; \Big[w\ \ log\ \Big(f\big(\{\boldsymbol{Z}_g^r\}\big|\ h_r = c\big)\Big)$$

$$+\ (1 - w)\big\{\ log\ \big(f(\{X_t\!: t \in W_r\}|h_r = c, \{n_t\!: t \in W_r\})\big)\ +\ log(P[h_r = c])\big\}\Big]$$

The term $f\big(\{\boldsymbol{Z}_g^r\}|\ h_r =\ c\big)$ reflects contribution from gene expression, while $f(\{X_t\!: t \in W_r\}|h_r = c, \{n_t\!: t \in W_r\})$ reflects contribution from SNPs.

We first focus on the latter term. It can be easily computed in log-scale as outlined below.

$$log\ f(\{X_t\!: t \in W_r\}|h_r = c, \{n_t\!: t \in W_r\})$$

$$= \sum_{t: t \in W_r} log\ f(X_t|h_r = c, n_t)$$

$$= \sum_{t:t \in W_r} log\left(\sum_{i=0}^{1} f(X_t, I_t = i | h_r = c, n_t)\right)$$

$$= \sum_{t:t \in W_r} log\left(\sum_{i=0}^{1} f(X_t | h_r = c, I_t = i, n_t)\ P[I_t = i]\right)$$

Now, we describe the computational details for $f(\{\boldsymbol{Z}_g^r\}|\ h_r = \ c)$ which denotes the contribution from gene expression data that cannot distinguish among the different ASCN configurations generating same total copy number. Additionally, it is reasonable to assume that SRT gene expression data, in general, is too sparse to distinguish between two copy deletions and a single copy deletion, or between a single copy amplification and various multi-copy amplifications. As a result, we have the following relationships.

$$f\left(\boldsymbol{Z}_g^r \middle| h_r = (0,0)\right) = f\left(\boldsymbol{Z}_g^r \middle| h_r = (0,1)\right),$$

$$f\left(\boldsymbol{Z}_g^r \middle| h_r = (1,1)\right) = f\left(\boldsymbol{Z}_g^r \middle| h_r = (0,2)\right),$$

$$f\left(\boldsymbol{Z}_g^r \middle| h_r = (1,2)\right) = f\left(\boldsymbol{Z}_g^r \middle| h_r = (1,3)\right) = f\left(\boldsymbol{Z}_g^r \middle| h_r = (2,2)\right)$$

Next, it can be shown that the conditional distribution of normalized expressions $\boldsymbol{Z}_g^r$ given any CNV state is same as the conditional distribution of $\boldsymbol{Z}_g^r$ given any ASCN configuration corresponding that CNV state. We derive this relationship below considering deletion as an example.

$$f\left(\boldsymbol{Z}_g^r \middle| s_r = 1\right)$$

$$= \frac{1}{P[s_r = 1]}\{ f\left(\boldsymbol{Z}_g^r \middle| h_r = (0,0)\right) P[h_r = (0,0)]$$

$$+ f\left(\boldsymbol{Z}_g^r \middle| h_r = (0,1)\right) P[h_r = (0,1)] \}$$

$$= f\left(\boldsymbol{Z}_g^r \middle| h_r = (0,0)\right)$$

$$= f\left(\boldsymbol{Z}_g^r \middle| h_r = (0,1)\right)$$

Similarly, we can derive the following relationships for the other two CNV states.

$$f\left(\boldsymbol{Z}_g^r \middle| s_r = 2\right) = f\left(\boldsymbol{Z}_g^r \middle| h_r = (1,1)\right) = f\left(\boldsymbol{Z}_g^r \middle| h_r = (2,2)\right)$$

$$f\left(\boldsymbol{Z}_g^r \middle| s_r = 3\right) = f\left(\boldsymbol{Z}_g^r \middle| h_r = (1,2)\right) = f\left(\boldsymbol{Z}_g^r \middle| h_r = (1,3)\right) = f\left(\boldsymbol{Z}_g^r \middle| h_r = (2,2)\right)$$

Finally, $\{f\left(\boldsymbol{Z}_g^r \middle| s_r = j\right): j = 1,2,3\}$ can be computed after fitting gene expression based GMM (Note 3.1). Then, we have

$f\left(\{\boldsymbol{Z}_{\boldsymbol{g}}^{\boldsymbol{r}}\} \middle| s_r = j\right) = \prod_g f\left(\boldsymbol{Z}_g^r \middle| s_r = j\right)$, where the product is taken with respect to the bins in region $r$.

#### 3.4 Impact of using known SNPs while calling germline SNPs from SRT dataset

We used a panel of common SNPs (minor allele frequency > 0.05) from phase 3 of the1000 Genomes Project [3] for identifying germline SNPs from SRT datasets. In principle, the allele counts at germline SNPs could have been obtained both with and without using the panel of known SNPs. In this section, we examined the role of known SNPs while extracting germline SNPs from SRT datasets.

For this assessment, we only considered the samples from patient 4 and patient 6 of the cSCC study [4]. We were unable to identify germline variants from the colon cancer scWGS data [5] without using the panel, as the annotations for benign cells were unavailable. We used Strelka2 to obtain germline variants from the normal WES data (Supplementary Figure 4 A). We observed that almost 84.8% (range: 84.4-85.1%) of the variants were present in the panel of known SNPs (Supplementary Figure 4 C-F). The variants, that were not part of the panel of known SNPs, were also considered as ground truth in this case. For the matched Visium samples, we used Strelka2 and bcftools to obtain germline variants from the set of benign locations, and found that, on an average, 50.8% (range:

47.3-55.7%) of those were common to the panel of known SNPs (Supplementary Figure 4 B, G-J). We did not use cellsnp-lite as it was not suitable for calling variants where the calls were to be restricted within certain intervals (e.g., exonic regions specified through a BED file). The effect of not considering known SNPs was most evident in the observed precision levels (Supplementary Figure 4 L-M). The precisions for both homozygous and heterozygous SNPs dropped irrespective of the variant-caller. Specifically, we observed an average fall in precision of 41.2% (range: 33.9-47.6%) for homozygous SNPs, and an average fall of 44% (range: 36.8-52.7%) for heterozygous SNPs. Thus, the panel of known SNPs acted as a filter in the final selection step, and significantly improved the signal to noise ratio in the resulting set of SRT SNPs. This observation bolsters the practice of using a panel of known SNPs while obtaining germline SNPs from sc-RNAseq or SRT data, particularly for CNV-analysis [6–8]. The recalls, F1 scores and the concordance rates either dropped slightly or remained unchanged as compared to the results obtained with the panel of known SNPs (Supplementary Figure 4 K-L).

#### 3.5 Assessment of phased SRT SNPs

We further examined the quality of phased germline heterozygous SNPs that can be derived from SRT data using a population-based phasing technique. Specifically, in the context of identifying ASCNs from sc-RNAseq and SRT data, the combination of cellsnp-lite and Eagle2 [9] had been extensively used to obtain phased heterozygous SNPs [6–8]. First, bi-allelic germline SNPs were identified using cellsnp-lite with the same panel of known SNPs mentioned before. Then, the resulting heterozygous SNPs are phased using Eagle2 based on the 1000 genomes phase 3 haplotypes. Different follow-up approaches had also been implemented to account for the phase-switch errors originating from such a population-based phasing technique (e.g., haplotype-aware HMM in [6]). Yet, the initial set of phased SNPs are expected to be of reasonable quality, and to the best of our knowledge, there exists no such quality assessment study especially focusing on SRT data.

With this motivation, we applied Eagle2 with the same reference panel on the heterozygous SNPs used as ground truth and on the heterozygous SNPs derived from SRT data using cellsnp-lite (Supplementary Figure 3 A, C). For SRT data, we used the pre-processing pipeline from Numbat [6] that combines cellsnp-lite and Eagle2 [6–8]. We defined a match between the two sets of phased heterozygous SNPs (ground truth and SRT) based on genomic location, allelic configuration and phasing configuration, and used precision and recall as evaluation criteria. Here, precision measured the percentage of matches within the SRT-derived calls and recall measured the percentage of matches among the true calls. The results revealed that the average precision was 47.3% (range: 41.8-51%), while average recall was 3.5% (range: 2.5-5%) (Supplementary Figure 3 D). We also noticed a platform-effect as opposed to the results discussed earlier without considering the phasing step. Specifically, the precision was higher in the two Visium samples compared to the Slide-seqV2 sample (avg. P= 50% vs. 41.8%). On the other hand, the recall for the Slide-seqV2 sample, although being poor, was almost twice of that for the Visium samples (avg. R= 2.7% vs. 5%). Interestingly, we had observed signal density (precision) of at least 90%, when we compared the locations of heterozygous SNPs obtained from SRT data against the locations of the corresponding ground truth (Main text). Furthermore, all the heterozygous SNPs with a location-match also had matched allelic configuration. But, the phased heterozygous SNPs derived using cellsnp-lite and Eagle2 had only about 47% signal density which might reflect the additional noise induced by phasing. The recall rates were consistently poor throughout our assessment. To summarize, we found that quality of the phased SNPs was not satisfactory, and might warrant further extensive examination that is beyond the scope of the current study. Additionally, future work can include quantification of the improvement induced by different approaches correcting for phase-switch errors especially in SRT datasets.

##### 3.6 ASCN estimation from colon cancer study

We applied Alleloscope [10] to infer ASCNs from scWGS data associated with the colon cancer study [5] (Supplementary Figure 12). Alleloscope generated a matrix (cell x chromosome) with estimated ASCNs. We examined ASCNs of the regions which were reported to harbor CNVs in the study [5]. These regions served as true CNVs in our analysis (deletion: 15p, 15q, 18p, and 18q; amplification: 1q, 7p, 7q, 8q, 20p, and 20q). Additionally, we considered ASCNs prevalent in at-least 25% cells.

The estimated ASCNs for chromosome 1 revealed that 78% cells had no CNVs (ASCN= (1,1)), while 19% cells harbored one copy amplification

(ASCN= (1,2)). On chromosome 7, 47% cells had no CNVs, alongside 49% cells harboring one copy amplification. Chromosome 20 also had a similar CNV profile with 36% cells having no CNVs and 42% cells harboring one copy amplification. Chromosome 8 had a different CNV profile. In addition to 28% cells with no CNVs, 43% cells had a total copy number of five with ASCN of (1,4), indicating three copy amplification of one haplotype chromosome. Notice that, such higher order amplifications are unlikely to be accurately inferred from SRT or sc-RNAseq datasets, and often are not considered in statistical models designed for such datasets [6]. Alleloscope did not generate results for chromosomes 15 and 18, the regions serving as true deletions in our analysis, as too few cells passed the quality control steps of the pipeline.

Next, we compared these results with the ASCNs estimated by SPICE from the colon cancer Slide-seqV2 dataset (Supplementary Table 1). We particularly focused on the accurate detections of SPICE. We observed that, SPICE successfully estimated one copy amplifications on chromosomes 1q, 7p, 20p and 20q. On the other hand, SPICE, being unable to estimate higher order amplification states such as (1,4) by design, inaccurately predicted one copy amplification on chromosome 8q.

Among the correct deletion calls, SPICE estimated one copy deletion on both chromosomes 15q and 18q and estimated two copy deletion on chromosome 18p. These deletions were prevalent in both subclones. As mentioned earlier, we were not able to validate the estimated ASCNs on chromosomes 15 and 18 due to absence of ground truth. We were inquisitive especially regarding SPICE's estimation of two copy deletion on chromosome 18p. Accordingly, we examined the B-Allele Frequencies (BAFs) or proportion of alternative alleles for heterozygous SNPs (Supplementary Figure 13). First, we noticed that there were only six heterozygous SNPs on chromosome 18p, reflecting extremely sparse SNP-density. Then, we found that all BAFs were zero in subclone 1 while five of the six BAFs were zero in subclone 2, validating SPICE's estimation of two copy deletion. Such extremes patterns were not observed in the BAFs of chromosome 18q, and accordingly the exploratory analysis supported SPICE's estimation of one copy deletion.

#### 3.7 ASCN estimation from cutaneous squamous cell carcinoma study

We used FACETS [11] to infer ASCNs from WES data generated from the tumor and its adjacent normal skin samples of patient 6 [4]. Our objective was to estimate ASCNs corresponding to the regions serving as true CNVs in our analysis (deletion: 3p, 4q, 5q, 10p, 10q, 13p, 13q, 21p, and 21q; amplification: 1p, 3q, 8q, 9q, 11q, 14q, 17q, 20p, and 20q). While examining the outputs from the pipeline, we excluded segments where only the total copy numbers were inferred, or segments spanning less than 1MB (Supplementary Figure 14, Supplementary Table 3) [11]. FACETS identified at least one genomic segment harboring amplification on each of the chromosome arms serving as true amplifications. On the contrary, it could not detect CNVs on the arms serving as true deletions except for inferring one copy deletions on chromosomes 3p and 21.

Next, we compared the ASCNs estimated by FACETS with the ASCNs estimated by SPICE from the patient 6 Visium dataset (Supplementary Table 2). SPICE was accurate in predicting one copy amplification with most of its correct amplification calls (1p, 9q, 20p, and 20q). On chromosome 9q, FACETS detected a one copy amplification, spanning about 65 MB, along with another much smaller segment, spanning only 3.6

MB, with ASCN of (0,4). Additionally, FACETS identified two segments (84 MB and 11 MB) with ASCN of (0,3) and one segment (8 MB) with ASCN of (0,4) on chromosome 3q where SPICE inaccurately predicted one copy amplification. Note that, complex ASCNs such as (0,3) and (0,4), involving higher order amplifications simultaneously with deletion are unlikely to be accurately inferred from SRT data alone.

As discussed in the main text, SPICE's likelihood-based integration framework is flexible in assigning more weightage to a specific modality. We found that the number genes on each chromosome arm, the average total UMI count, and the average number of genes expressed at each location were higher in the Visium samples (Supplementary Figure 1 A-B, E-G). On the other hand, the average total read count per heterozygous SNP was lower on each arm for the Visium dataset of patient 6, reflecting lower confidence on the SNP modality (Supplementary Figure 2 B). We utilized this observation by assigning 25% additional weightage to SPICE's gene expression component ($w$=0.75; Note 3.3). As a result, there was a substantial boost in SPICE's performance as it additionally identified one copy deletion on chromosome 3p and one copy amplification on chromosome 8q.

#### 3.8 Sensitivity analysis of SPICE

We conducted a sensitivity analysis of SPICE's performance considering two aspects: the weighting scheme used to integrate the two data modalities into a single joint likelihood framework (Note 3.3), and the weighting scheme associated with the prior probabilities of different ASCN states (Note 3.2). To assess the sensitivity with respect to the weights used in integration, we selected four different weights: $w = 0.35$, 0.50, 0.75, and 0.85. Our default choice of $w = 0.50$ assigned equal weightage to each modality. Notice that, $w = 0.35$ sets 35% weight to gene expression component, while $w = 0.85$ assigns a high weightage of 85% to it. We had observed that the GMM applied to gene expression had an edge in identifying true CNVs, while the Binomial mixture model, used for SNPs, primarily provided safeguard against false detections. Accordingly, a higher weightage to the gene expression component is expected to generate higher recalls, whereas a higher weightage to the SNP component is likely to benefit SPICE with a higher precision.

To assess the sensitivity with respect to the weights associated with the prior probabilities of ASCNs, we have selected four different choices (0.60, 0.90, 0.95, 0.99) for each of $\zeta_{01}$ and $\zeta_{12}$. Recall that $\zeta_{01}$ reflects the weightage of the one copy deletion state (i.e., ASCN= (0,1)), between the two ASCN states related to deletion (i.e., ASCNs (0,0) and (0,1)). Thus, $\zeta_{01}$>0.50 assigns higher prior probability on one copy deletion between the two deletion states. Similarly, $\zeta_{12}$ reflects the weightage of one copy amplification state (i.e., ASCN= (1,2)) among the ASCN states reflecting amplification (i.e., ASCNs (1,2), (1,3) and (2,2)). Our choices for the sensitivity analysis reflect the prior belief that one copy deletion or one copy amplification are the most likely CNV-harboring ASCNs that can be accurately inferred from SRT data.

The results (Supplementary Figure 9- Supplementary Figure 11) illustrate SPICE's ability to generate stable and robust results. We first discuss results from the colon cancer Slide-seqV2 dataset (Supplementary Figure 9). SPICE's recall remained unchanged across all configurations (R=80%; 64 possible configurations of $w$, $\zeta_{01}$, and $\zeta_{12}$). But, SPICE's precision gradually dropped with decreasing weight to the SNP modality (avg. P=65%; range=50-80%), and consequently, the F1 score also decreased (avg. F1=71.5%; range=61.5-80%). These findings were consistent with the results discussed in the main text in the sense that the SNP component of SPICE provided a safeguard against false CNV calls. Notice that the minimum F1 score observed was still the highest considering the F1 scores achieved by the competing methods. Next, we explicitly focused on the default weighting scheme used in SPICE (i.e., $w = 0.50$), and examined the results with respect to changes in $\zeta_{01}$ and $\zeta_{12}$. First, we observed less variation in the results (avg. P=62.8% with range=61.5-66.7%; avg. F1=70.4% with range=69.6-72.7%), reflecting higher stability. Second, we noticed a minor drop (~4%) in average F1 score as $\zeta_{01}$ increased, while the results did not change on an average with respect to $\zeta_{12}$.

In the case of patient 6 Visium dataset (Supplementary Figure 10), precision showed a decreasing trend while recall had an increasing trend as more weight was assigned to gene expression. The F1 score also increased alongside recall, and the best F1 score was observed with least weightage on the SNPs. Here, the recall of SPICE was adversely impacted by the poor recall of the SNP component which provided perfect precision with only three CNV calls. Thus, SPICE's performance improved as more weightage was assigned to the gene expression component. Next, we considered the equal weighting scheme ($w = 0.50$), to examine changes with respect to $\zeta_{01}$ and $\zeta_{12}$. The results remained largely unchanged on an average with respect to $\zeta_{01}$. The recall and the F1 score gradually dropped on an average as $\zeta_{12}$ increased (range for avg. R= 27.8-34.7%; range for avg. F1= 41.7-49.5%), while the precision levels had an increasing trend with respect to $\zeta_{12}$ (range for avg. P= 83.3-100%).

Finally, for patient 4 Visium dataset (Supplementary Figure 11), SPICE achieved zero false detection across all configurations.

#### 3.9 Additional details on computational tools

##### Eagle2

We used Eagle2 (version 2.4.1) for phasing bi-allelic germline heterozygous SNPs based on the publicly available 1000 genomes phase 3 haplotypes as a reference panel. This reference panel was downloaded from Numbat's website [6]. We used the hg38 based panel for the cSCC datasets (WES and Visium), and the hg19 based panel for the colon cancer datasets (scWGS and Slide-seqV2).

##### Alleloscope

We applied Alleloscope (version 1.0.1) [10] to infer ASCNs from scWGS data associated with the colon cancer study [5]. We used chromosomes as segments, and followed steps 0-6 from the tutorial https://github.com/seasoncloud/Alleloscope/tree/main/samples/P5931/scDNA. Specifically, we used cellsnp-lite [12] with a panel of known SNPs (common SNPs from phase 3 of the1000 Genomes Project; see ‘Panel of known SNPs’ in the main text for further details) to generate the required VCF file, and used VarTrix (version 1.1.22; https://github.com/10XGenomics/vartrix) to generate the allele-count matrices.

**FACETS**

We used FACETS [11] to infer ASCNs from WES data generated from the tumor and its adjacent normal skin samples of patient 6 [4]. We followed cnv_facets: https://github.com/dariober/cnv_facets (version 0.16.0) for implementation.

We specified the exonic regions through the Agilent target BED file (SureSelect Human All Exon V6 r2; hg38), and used a panel of known SNPs (common SNPs from phase 3 of the1000 Genomes Project; see ‘Panel of known SNPs’ in the main text for further details) as the required VCF file.

### 4. Supplementary Figures

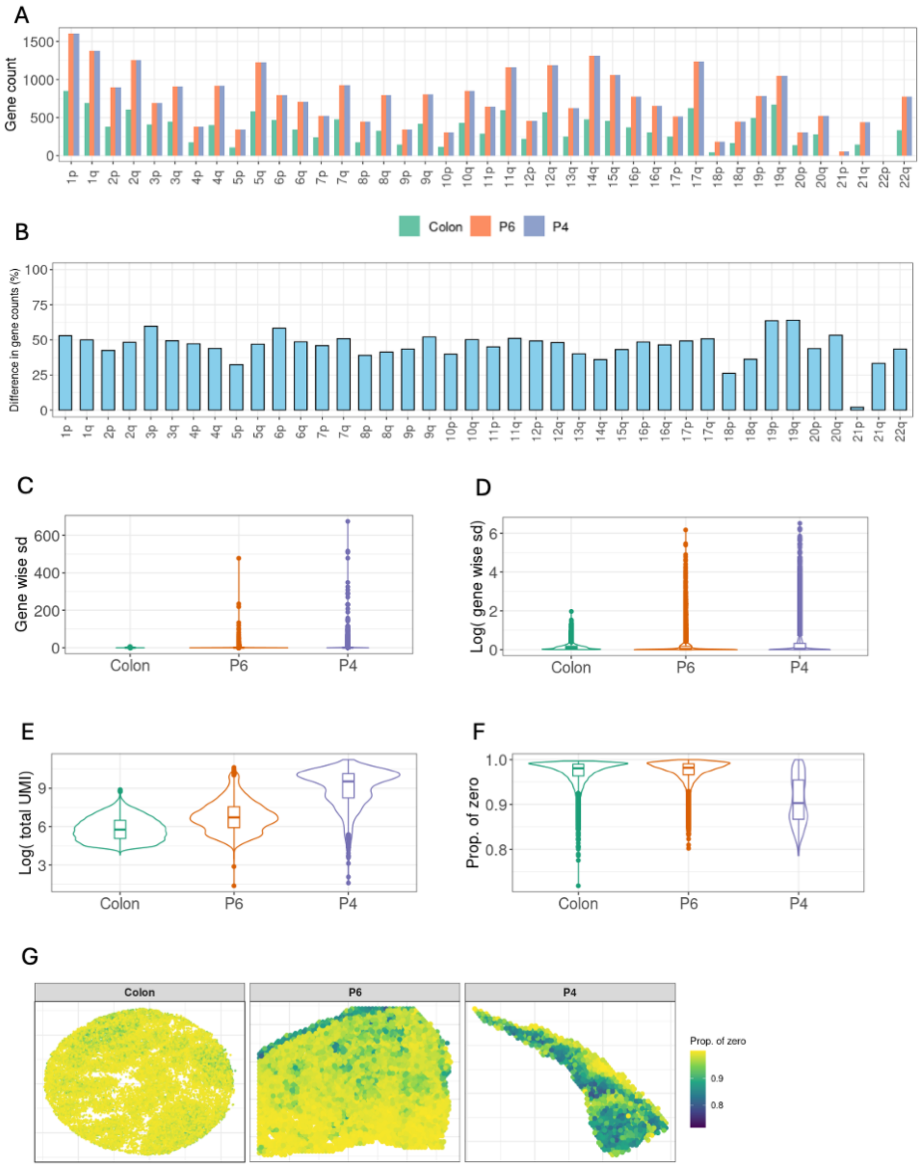

**Supplementary Figure 1. Features of SRT gene expression data.**
**A**. Number of mapped genes on each chromosome arm. The two Visium samples (P6, P4) have identical number of genes on each arm. **B**. Percentage difference in number of genes on each arm for colon cancer Slide-seqV2 dataset as compared to either of the two Visium datasets shown in A. **C-F**. Violin plots and boxplots of different features. Embedded boxplots indicate the median (center line) and interquartile range/IQR (box). Points represent outliers lying beyond whiskers (1.5x IQR). The features shown are, standard deviation (sd) of UMI counts for each gene (C), same as C in log-scale (D), log transformed total UMI counts for each location (E), and proportion of genes with zero expression for each location (F). **G**. Same as F shown over the SRT tissue locations. Autosomal genes are considered. Colon: colon cancer study, P4: patient 4 from cSCC study, P6: patient 6 from cSCC study.

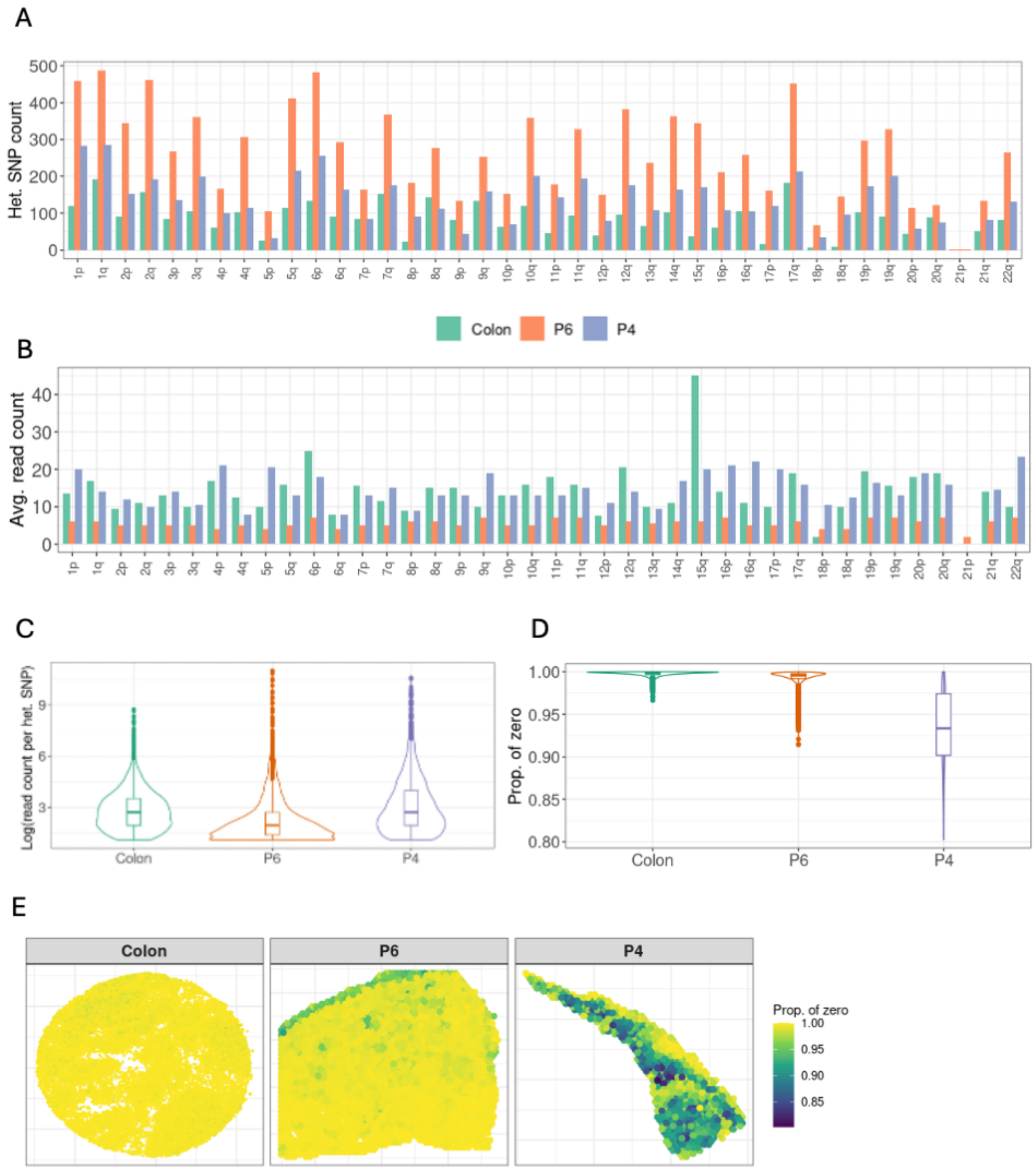

**Supplementary Figure 2. Features of heterozygous SNPs obtained from SRT data.**
**A**. Number of heterozygous SNPs on each chromosome arm. cellsnp-lite was used for SNP calling.
**B**. Average (median) read count of heterozygous SNPs on each chromosome arm. Read count refers to sum of the number of reads mapped to the two alleles of a heterozygous SNP. Median was taken over heterozygous SNPs mapped to each arm. **C-D**. Violin plots and boxplots of different features. Embedded boxplots indicate the median (center line) and interquartile range/IQR (box). Points represent outliers lying beyond whiskers (1.5x IQR). The features shown are, log transformed total read count for each heterozygous SNP (C) and proportion of heterozygous SNPs with zero total read count for each location (D). **E**. Same as D shown over the SRT tissue locations. Autosomes are considered. Colon: colon cancer study, P4: patient 4 from cSCC study, P6: patient 6 from cSCC study.

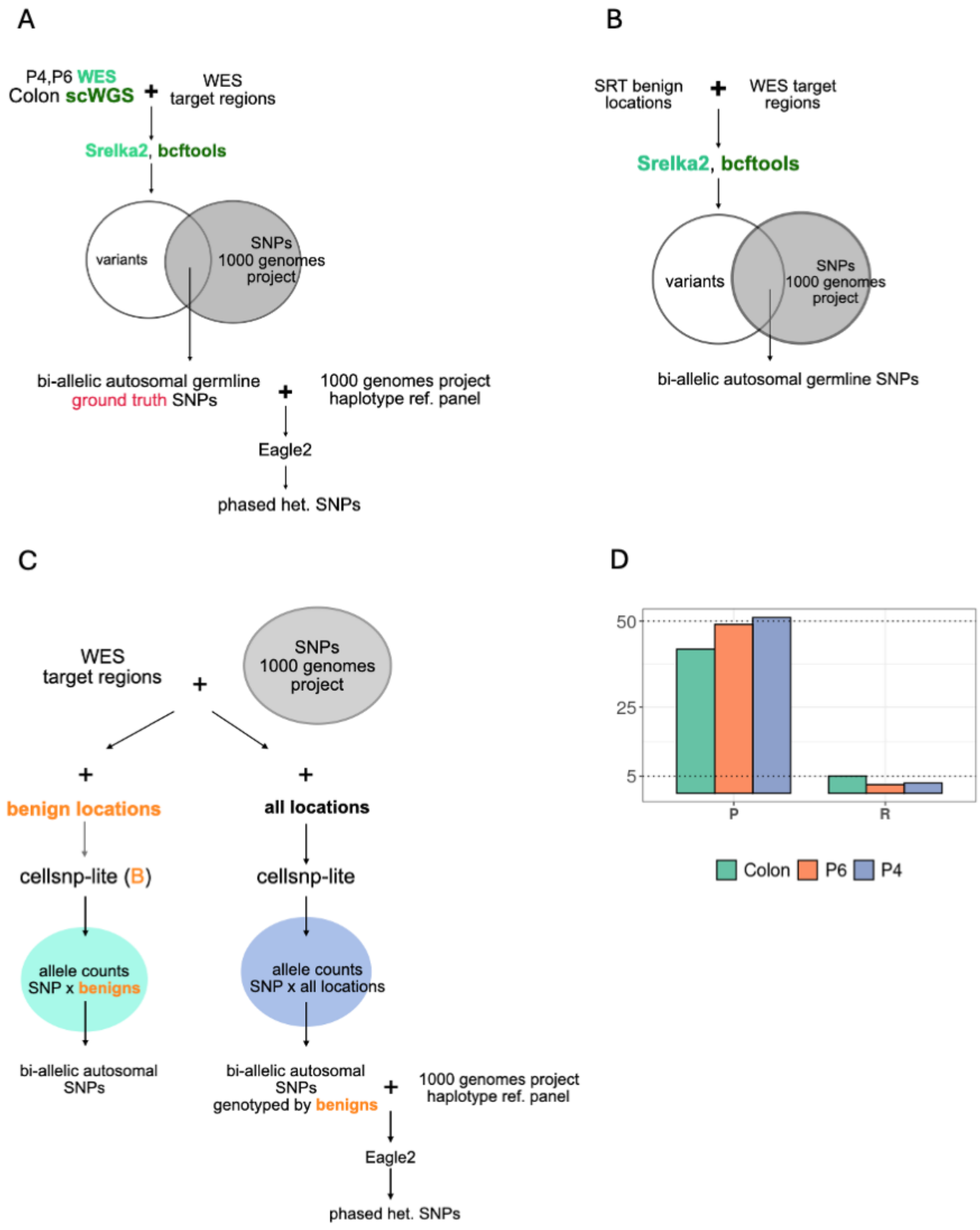

**Supplementary Figure 3. Schematics for germline SNPs calling.**
**A.** Schematic showing steps involved in calling germline SNPs from WES and scWGS data. **B-C.** Schematics showing steps to call germline SNPs from SRT data. Steps involved while using Strelka2 and bcftools (B). Steps involved with cellsnp-lite (C). We used two different approaches with cellsnp-lite. 'cellsnp-lite (B)' denotes the approach where a bam file having only the benign locations was used. 'cellsnp-lite' denotes the approach where the original bam file was used. **D.** Precision and recall (in percentages) for assessing phased heterozygous SNPs. Horizontally dotted lines mark 10 and 50. Colon: colon cancer study, P4: patient 4 from cSCC study, P6: patient 6 from cSCC study.

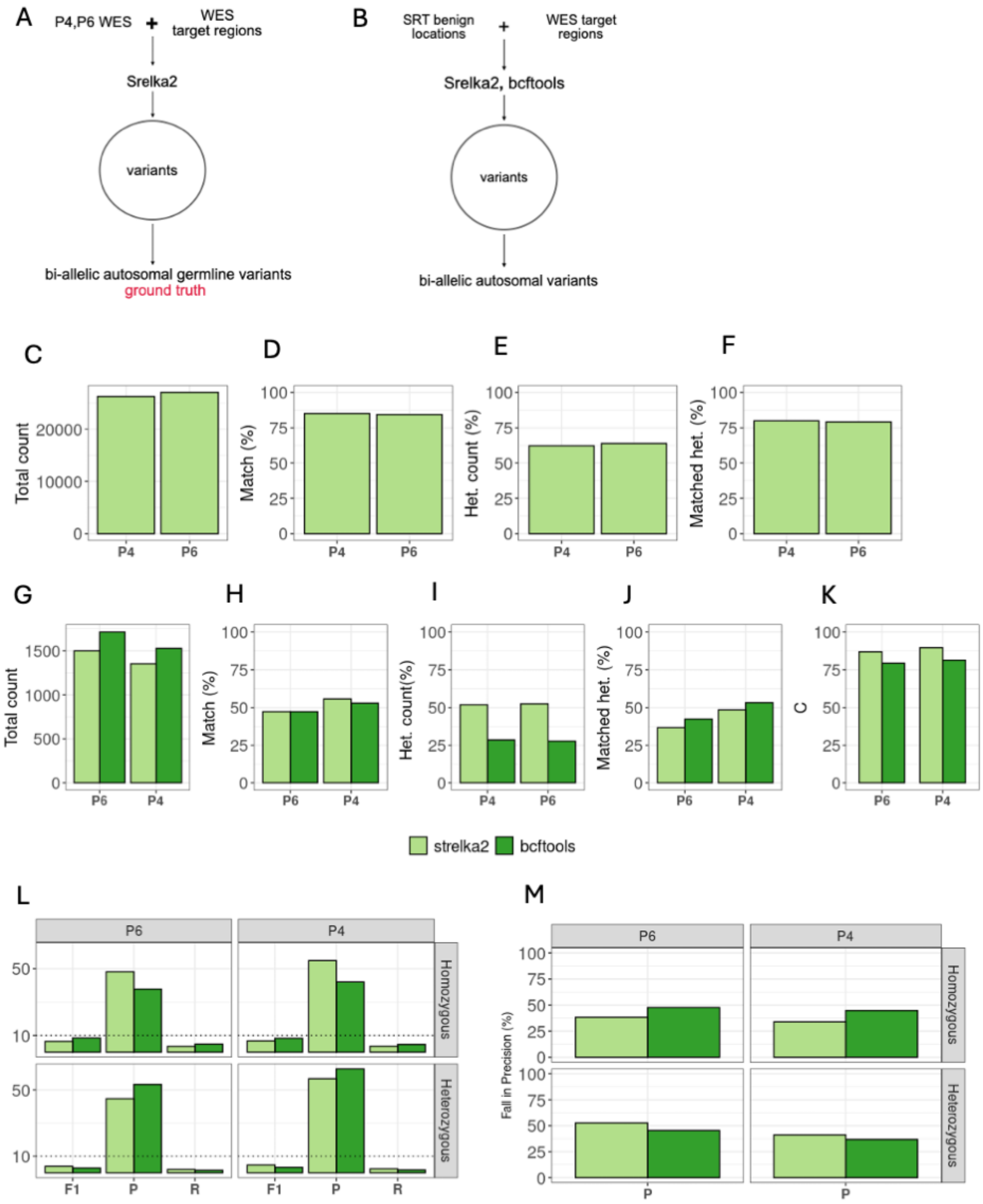

**Supplementary Figure 4. Quality assessment of variants called from SRT samples without using known SNPs.**
**A-B**. Schematics showing steps used for germline variant calling from WES (A) and SRT samples (B). **C-F**. Details on germline variants obtained from WES samples. Total variant count (C), percentage of variants present in the panel of known SNPs (D), percentage of heterozygous variants (E), and percentage of heterozygous variants present in the panel of known SNPs (F). **G-K.** Details on germline variants obtained from SRT samples. Total variant count (G), percentage of variants, combining heterozygous and homozygous, present in the panel of known SNPs (H), percentage of heterozygous variants (I), percentage of heterozygous variants present in the panel of known SNPs (J), and concordance rates (K). **L.** F1 score, precision and recall (in percentage). Horizontally dotted line marks 10. **M.** Percentage drop in precision when the panel of known SNPs is not used.
P4: patient 4 from cSCC study, P6: patient 6 from cSCC study.

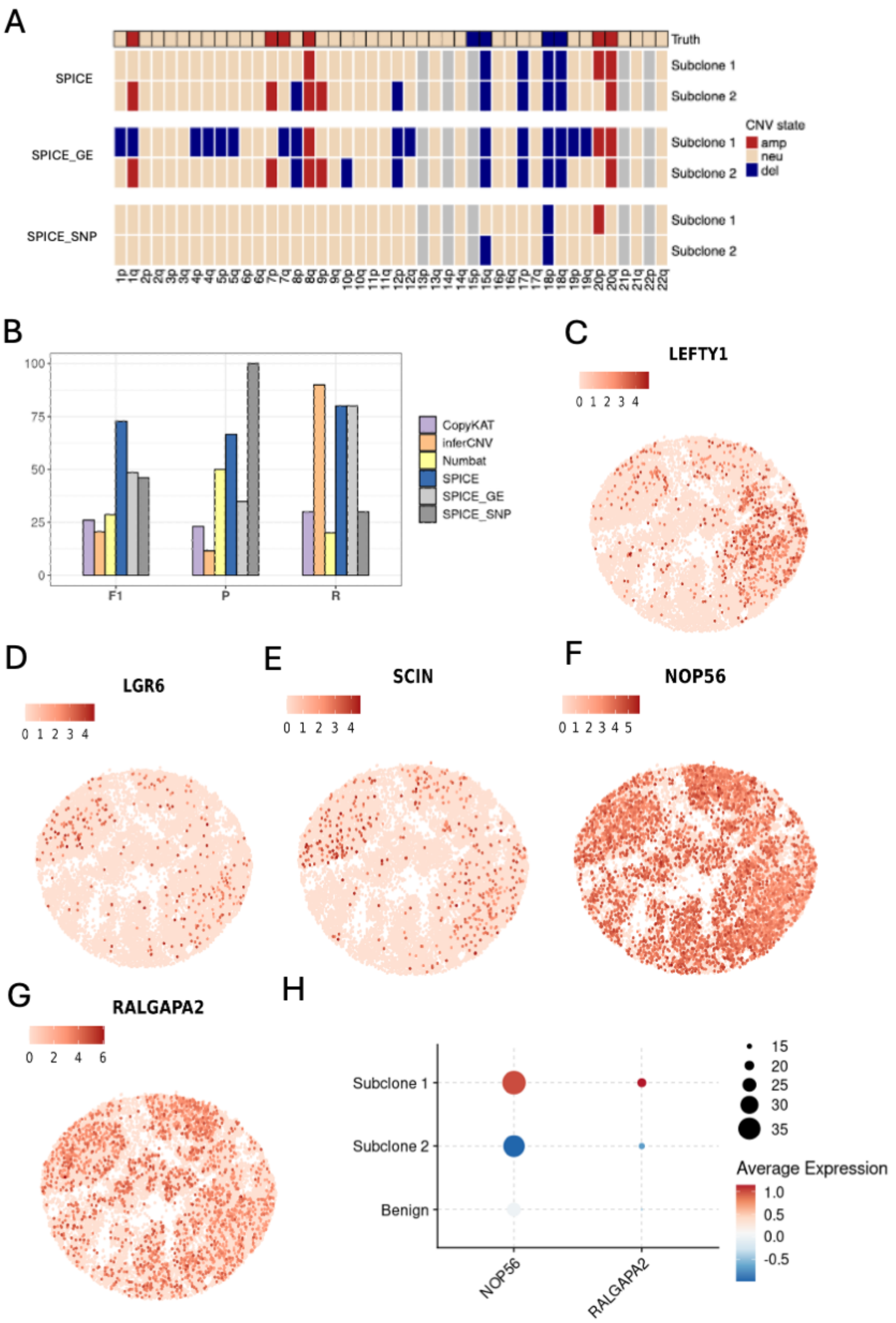

**Supplementary Figure 5. Additional results on CNV-detection in colon cancer Slide-seqV2 dataset.**
**A**. CNV states estimated by SPICE and its two components: gene expression component (SPICE _GE) and SNP component (SPICE _SNP), within each subclone for each chromosome arm. True CNV states are shown in the top bar. Grey denotes cases where estimation was not possible due to insufficient data points. **B**. F1 score, precision and recall (in percentage) for SPICE, SPICE _GE, SPICE _SNP, and the competing methods. **C-G**. Examples of genes characterizing the two subclones identified by SPICE. Differential expression analysis was performed between the subclones, and DE genes were mapped to CNV harboring regions successfully traced by SPICE. Plotted genes had well-established roles in colorectal cancer biology. NOP56 (20p) and RALGAPA2 (20p) were upregulated in subclone 1, while LEFTY1 (1q), LGR6 (1q), and SCIN (7p) were upregulated in subclone 2. Normalized expressions are plotted (layer='data') using Seurat::FeaturePlot(). **H.** Dot plot visualization of NOP56 and RALGAPA2 over the begin region and the two subclones. These genes were upregulated in subclone 1 where arm 20p harbored subclonal amplification. The size of the dot encodes the percentage of locations within a class. Created with Seurat::DotPlot().

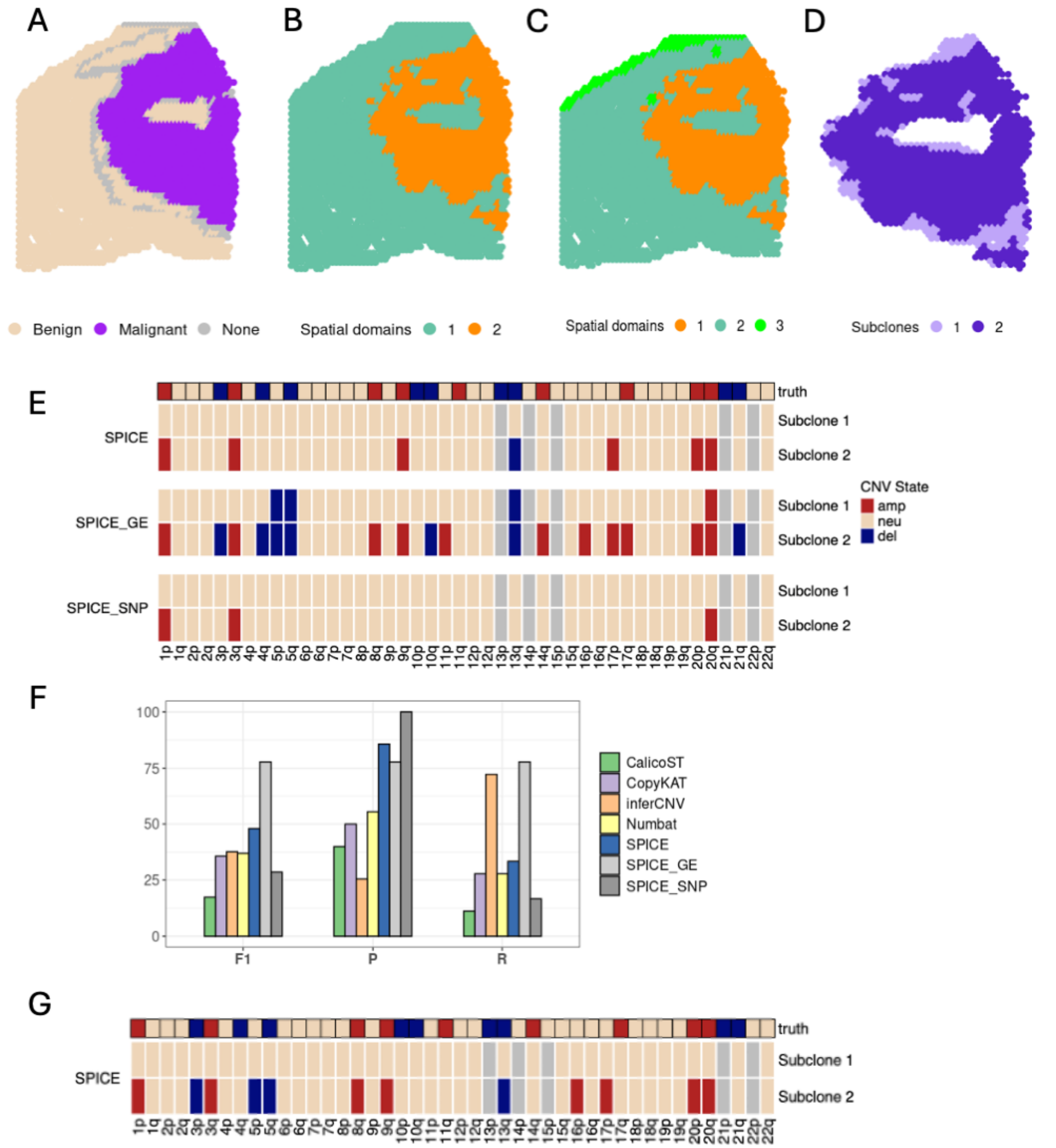

**Supplementary Figure 6. Additional results on CNV-detection in patient 6 Visium dataset.**
**A**. Scatter plot showing annotated locations. **B-D**. Scatter plot showing SpatialPCA-derived clusters (spatial domains) with number of clusters set as 2 (B) and 3 (C). In both cases, malignant locations, shown in A, were partitioned identically into two subclones (D). **E**. CNV states estimated by SPICE and its two components: gene expression component (SPICE _GE) and SNP component (SPICE _SNP), within each subclone for each chromosome arm. True CNV states are shown in the top bar. Grey denotes cases where estimation was not possible due of absence of data or too few data points. **F**. F1 score, precision and recall (in percentage) for SPICE, SPICE _GE, SPICE _SNP, and the competing methods. **G**. CNV states estimated by SPICE with 75% weightage to SPICE _GE. The legends are same as those in E.

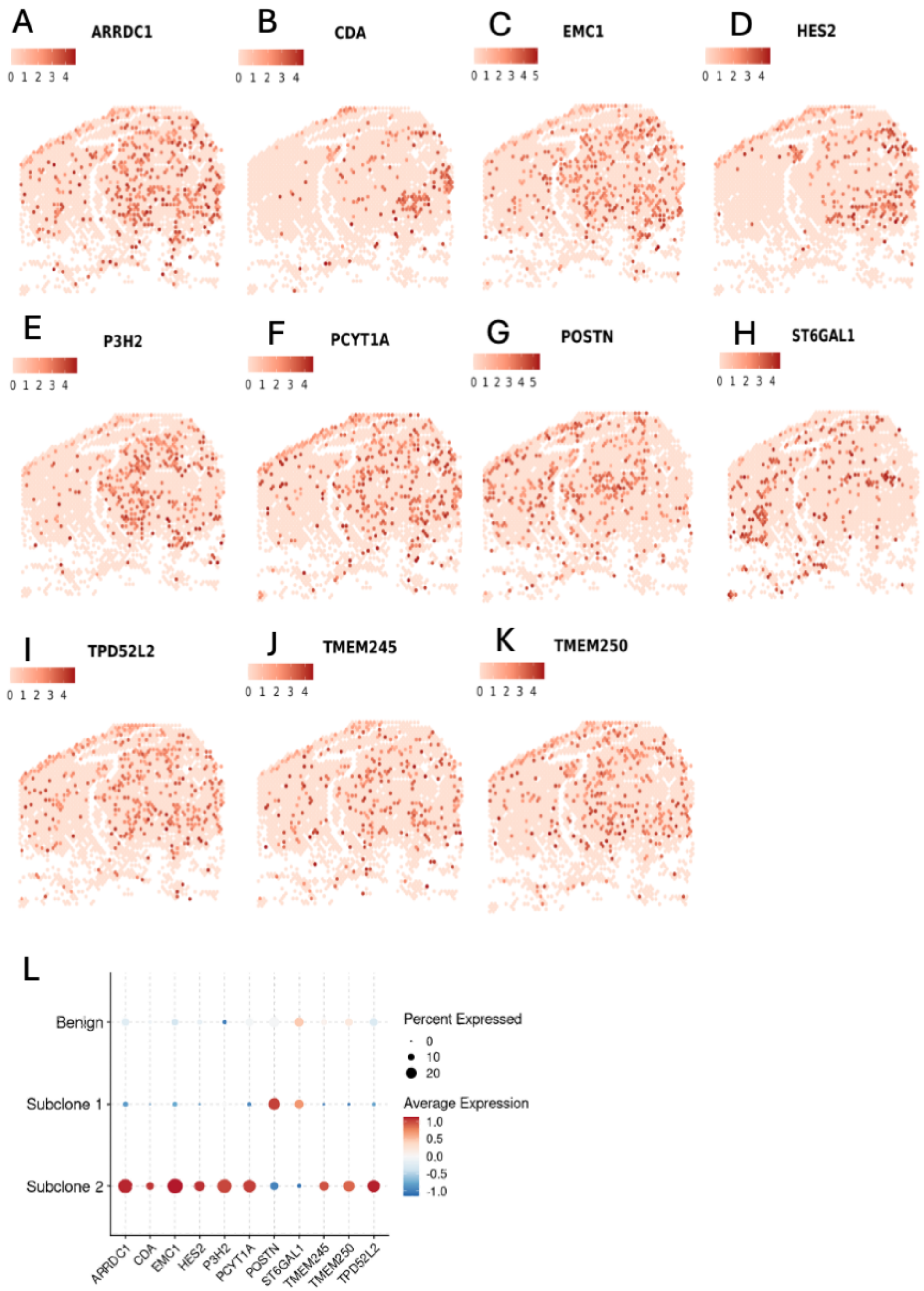

**Supplementary Figure 7. Gene expression characterizing subclones identified by SPICE in patient 6 Visium dataset.**

**A-K**. Examples of genes characterizing the two subclones identified by SPICE (Supplementary Figure 6 D). Differential expression analysis was performed between the two subclones, and DE genes were mapped to the CNV harboring regions successfully traced by SPICE (CDA, HES2, & EMC1 are on 1p. P3H2, ST6GAL1, & PCYT1A are on 3q. TMEM250, ARRDC1, & TMEM245 are on 9q. TPD52L2 is on 20q). POSTN (13q) showed reduced expression in subclone 2 where 13q had undergone deletion. The normalized expressions are plotted (layer='data') using Seurat::FeaturePlot(). **L**. Dot plot visualization of genes in A-K over the begin region and over the subclones identified by SPICE. The size of the dot encodes the percentage of locations within a class. Created using Seurat::DotPlot().

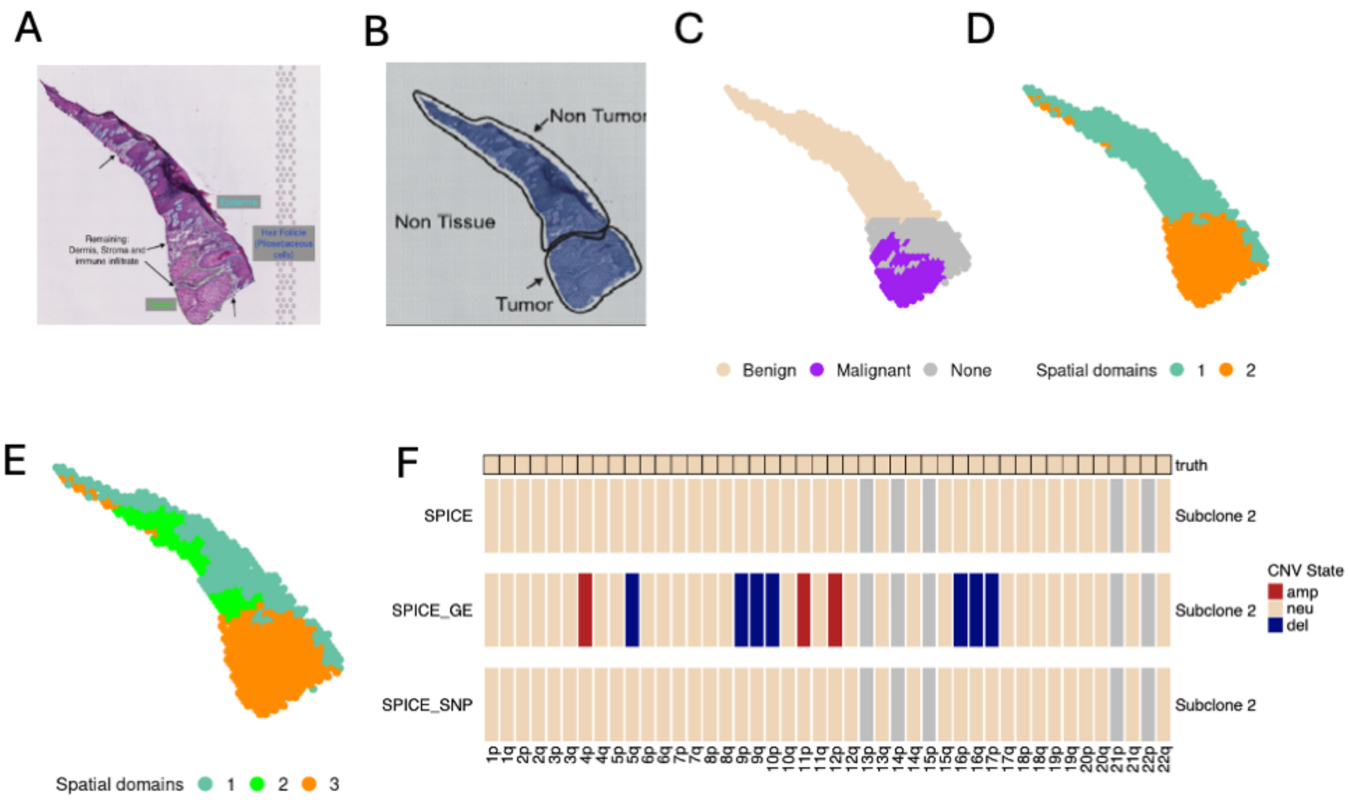

**Supplementary Figure 8. SPICE's performance in patient 4 Visium dataset.**
**A-B**. H&E images with annotations. A and B have respectively been extracted from Fig. S3 in [13] and Supp. Fig. 18A in [14]. **C**. Scatter plot showing annotated locations obtained using A and B. **D-E**. Scatter plot showing SpatialPCA-derived clusters (spatial domains) with number of clusters set as 2 (D) and 3 (E). Malignant locations, shown in C, belonged to a single spatial domain in both cases. **F**. CNV states estimated by SPICE and its two components: gene expression component (SPICE _GE) and SNP component (SPICE _SNP), within each subclone for each chromosome arm. Subclone 2 refers to the malignant locations belonging to spatial domain 2 in D. True CNV states are shown in the top bar. Grey denotes cases where estimation was not possible due of absence of data or too few data points.

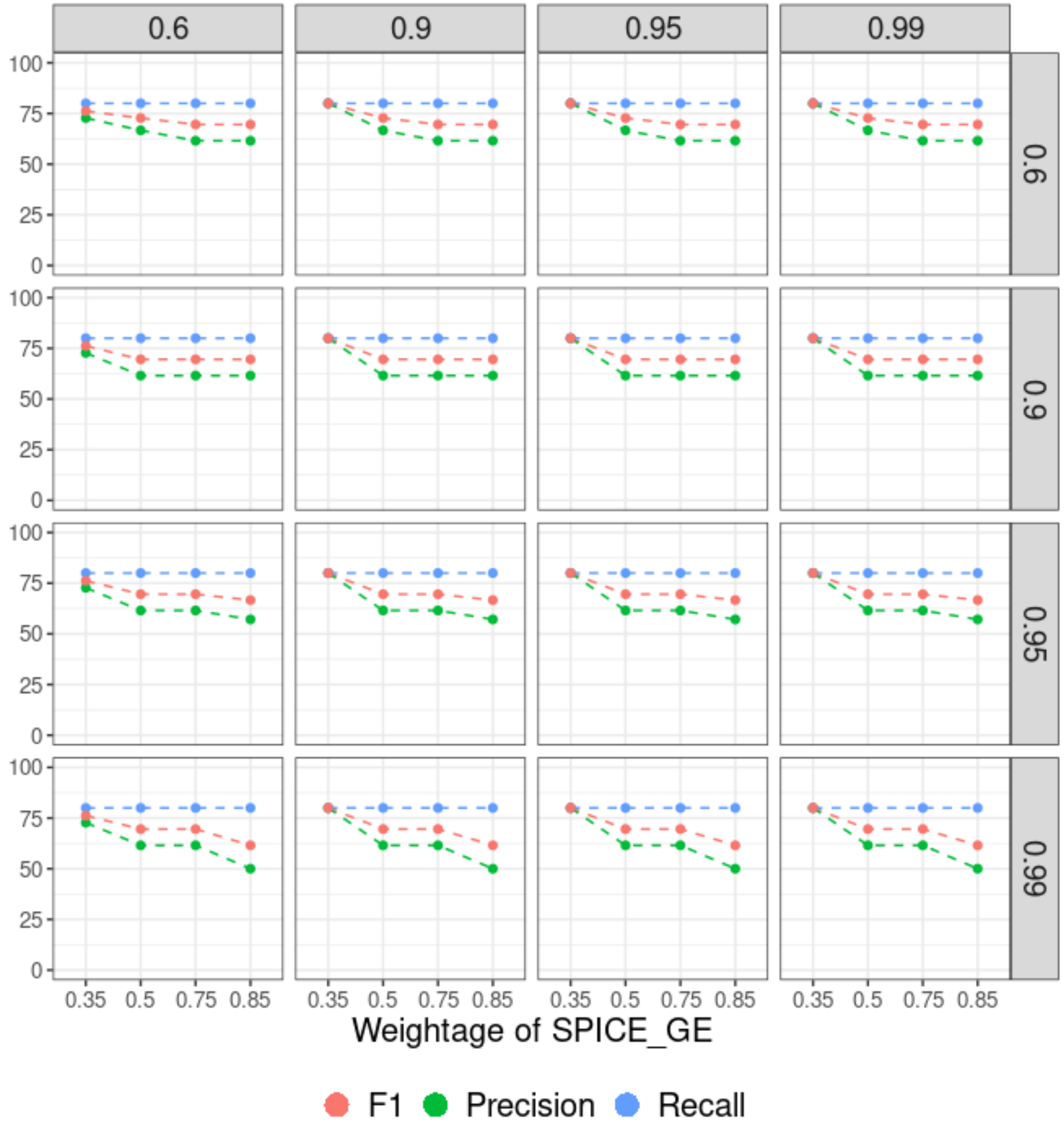

**Supplementary Figure 9. SPICE's performance across different weighting schemes for colon cancer Slide-seqV2 dataset.**
Precision, Recall, and F1 score of SPICE across different weighting schemes for colon cancer Slide-seqV2 dataset. Fractional weights ($w$; Note 3.3) assigned to the gene expression component of SPICE (SPICE_GE) are shown along the x-axis. Fractional weights corresponding to the prior probability of one copy deletion ($\zeta_{01}$; Note 3.2) are shown along the rows. Fractional weights corresponding to the prior probability of one copy amplification, ($\zeta_{12}$; Note 3.2) are shown along the columns.

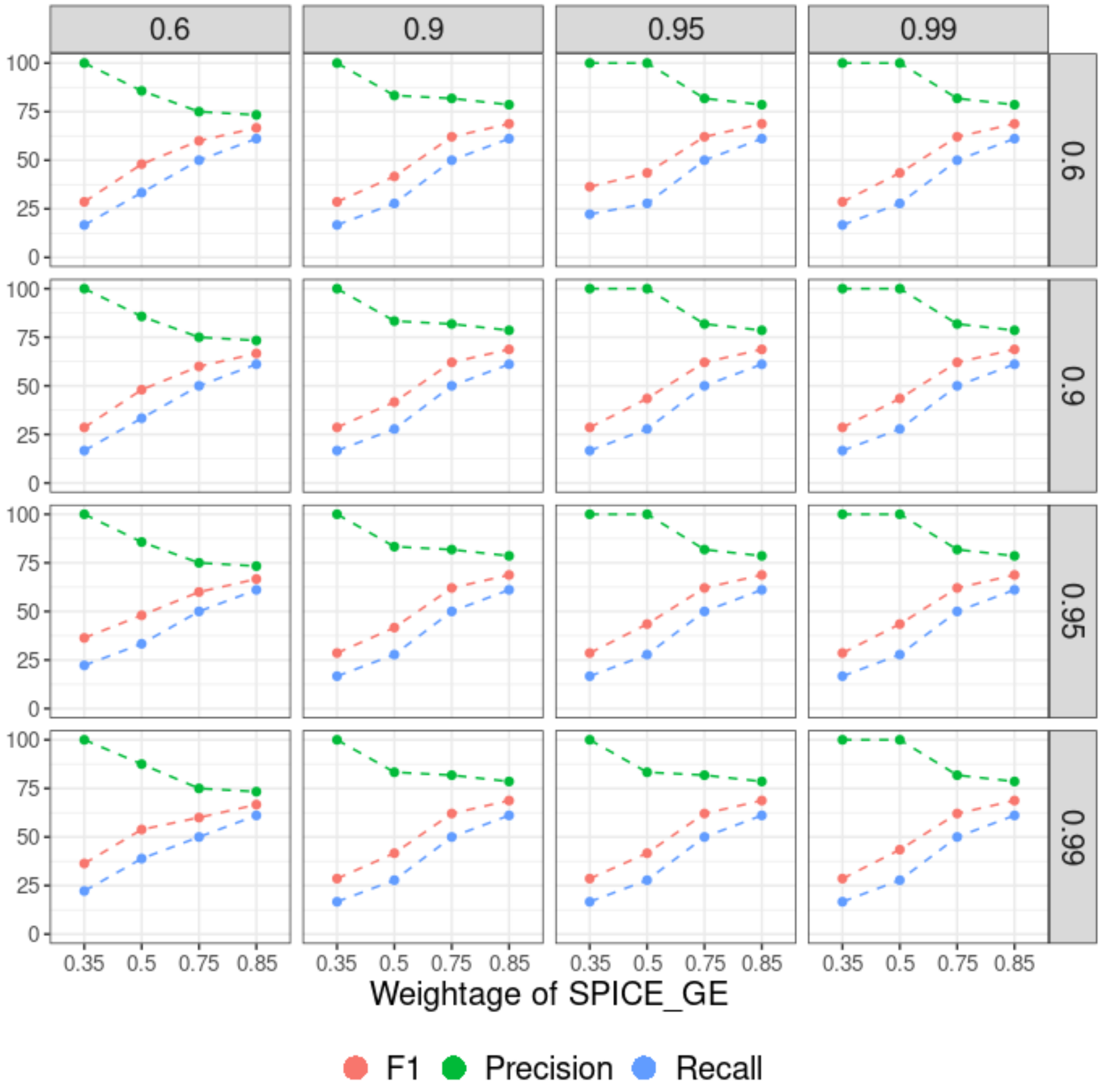

**Supplementary Figure 10. SPICE's performance across different weighting schemes for patient 6 Visium dataset.**

Precision, Recall, and F1 score of SPICE across different weighting schemes for patient 6 Visium dataset. Fractional weights ($w$; Note 3.3) assigned to the gene expression component of SPICE (SPICE_GE) are shown along the x-axis. Fractional weights corresponding to the prior probability of one copy deletion ($\zeta_{01}$; Note 3.2) are shown along the rows. Fractional weights corresponding to the prior probability of one copy amplification, ($\zeta_{12}$; Note 3.2) are shown along the columns.

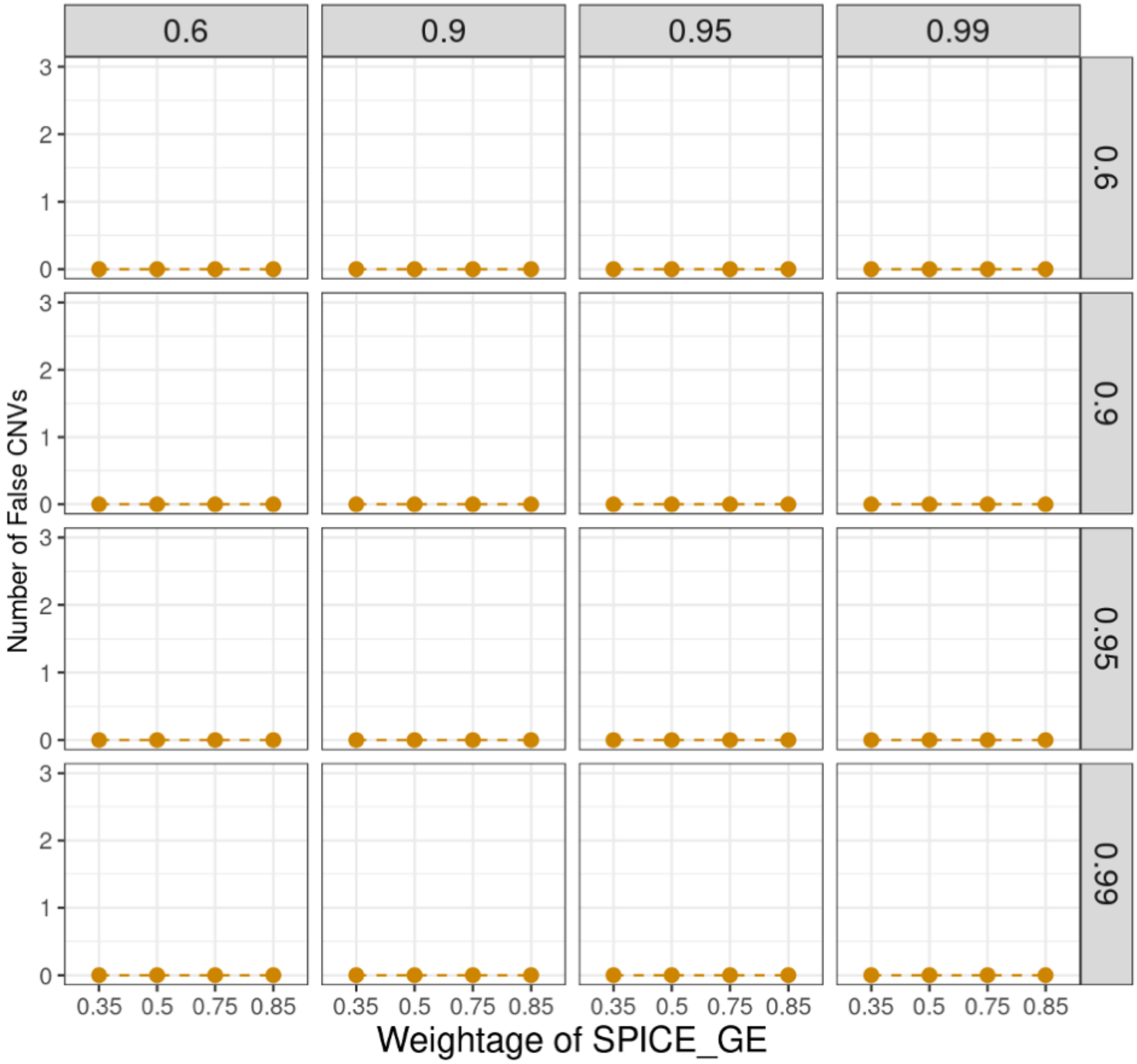

**Supplementary Figure 11. SPICE's performance across different weighting schemes for patient 4 Visium dataset.**
Number of false CNVs identified by SPICE across different weighting schemes for patient 4 Visium dataset. Fractional weights ($w$; Note 3.3) assigned to the gene expression component of SPICE (SPICE_GE) are shown along the x-axis. Fractional weights corresponding to the prior probability of one copy deletion ($\zeta_{01}$; Note 3.2) are shown along the rows. Fractional weights corresponding to the prior probability of one copy amplification, ($\zeta_{12}$; Note 3.2) are shown along the columns.

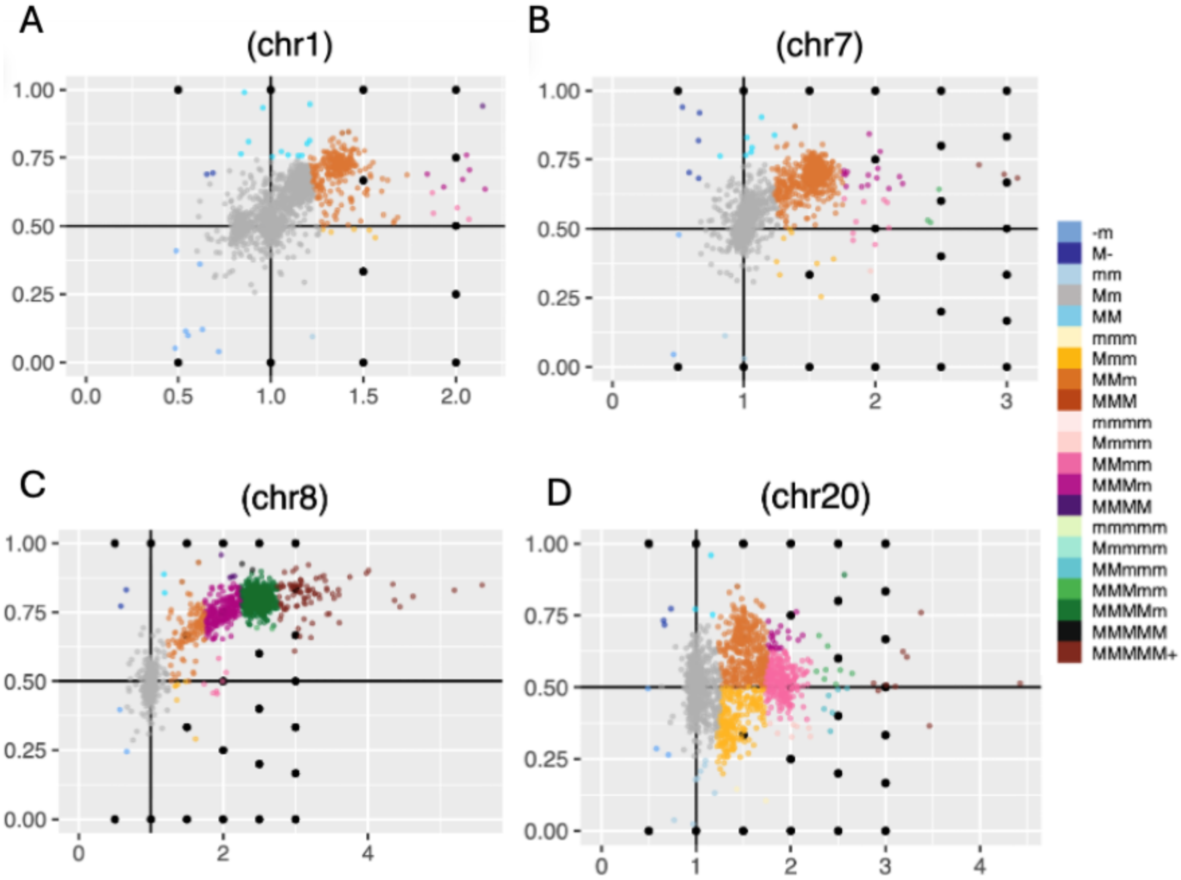

**Supplementary Figure 12. ASCN estimation from colon cancer scWGS dataset using Alleloscope. A-D**. Genotyping results, generated by Alleloscope, for chromosomes serving as true CNVs. The color scheme and legend bar are same as those used in Fig. 2b of [10]. The haplotype chromosome with higher number of copies across cells is called the 'major haplotype' (M), and the other one is called the 'minor haplotype' (m). Refer to [10] for further details. Alleloscope did not generate results for chromosomes 15 and 18, the regions serving as true deletions, as too few cells passed the quality control.

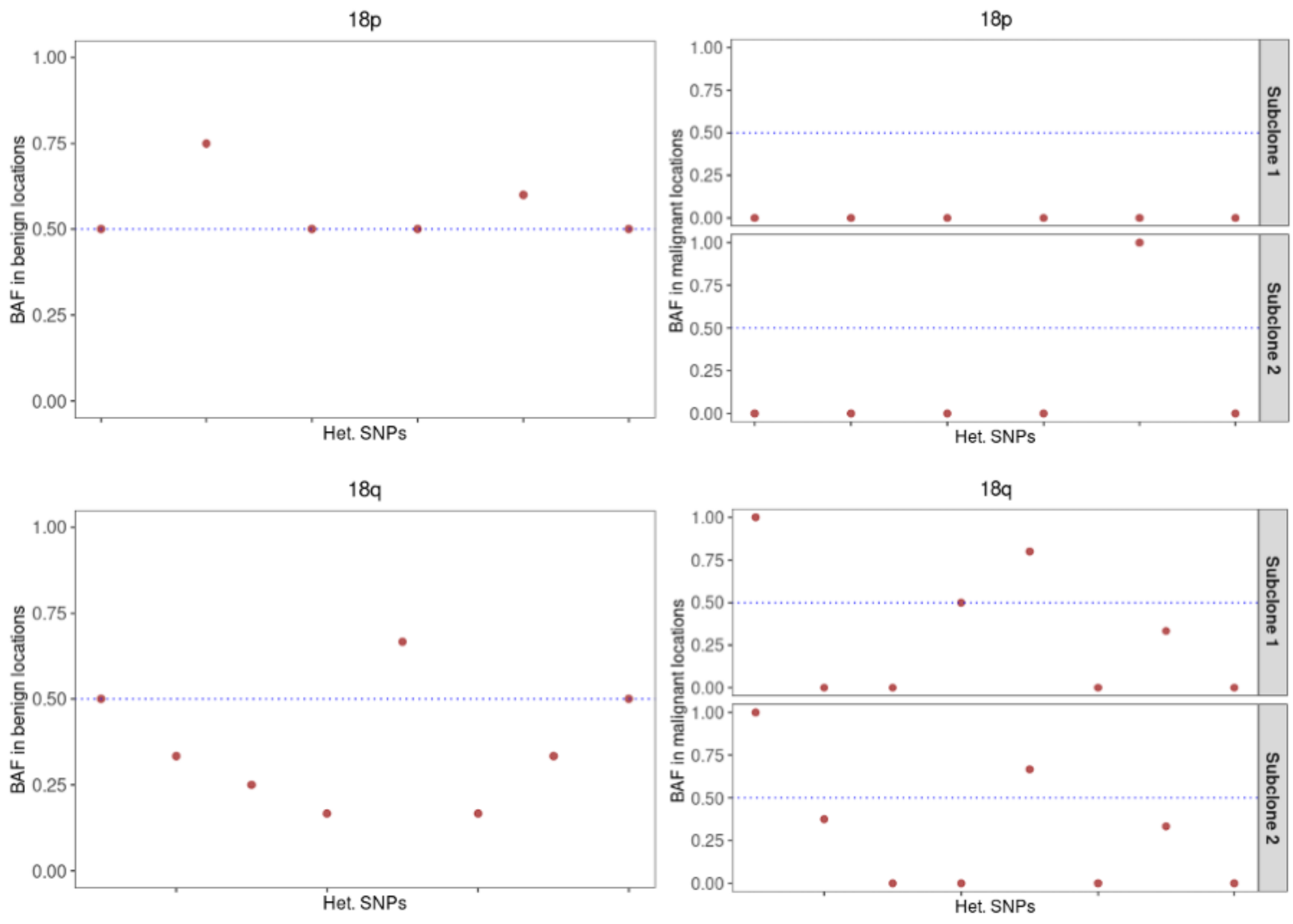

**Supplementary Figure 13. B-allele frequency plots on chromosome 18 from colon cancer Slide-seqV2 dataset.**
B-Allele Frequencies (BAFs) or proportion of alternative alleles are plotted for heterozygous SNPs on chromosome 18p (top panel) and on chromosome 18q (bottom panel). B-Allele frequencies computed from benign locations (left column) and from malignant subclones (right column) are shown separately. A horizontal line is drawn at 0.5, the ideal BAF in benign region.

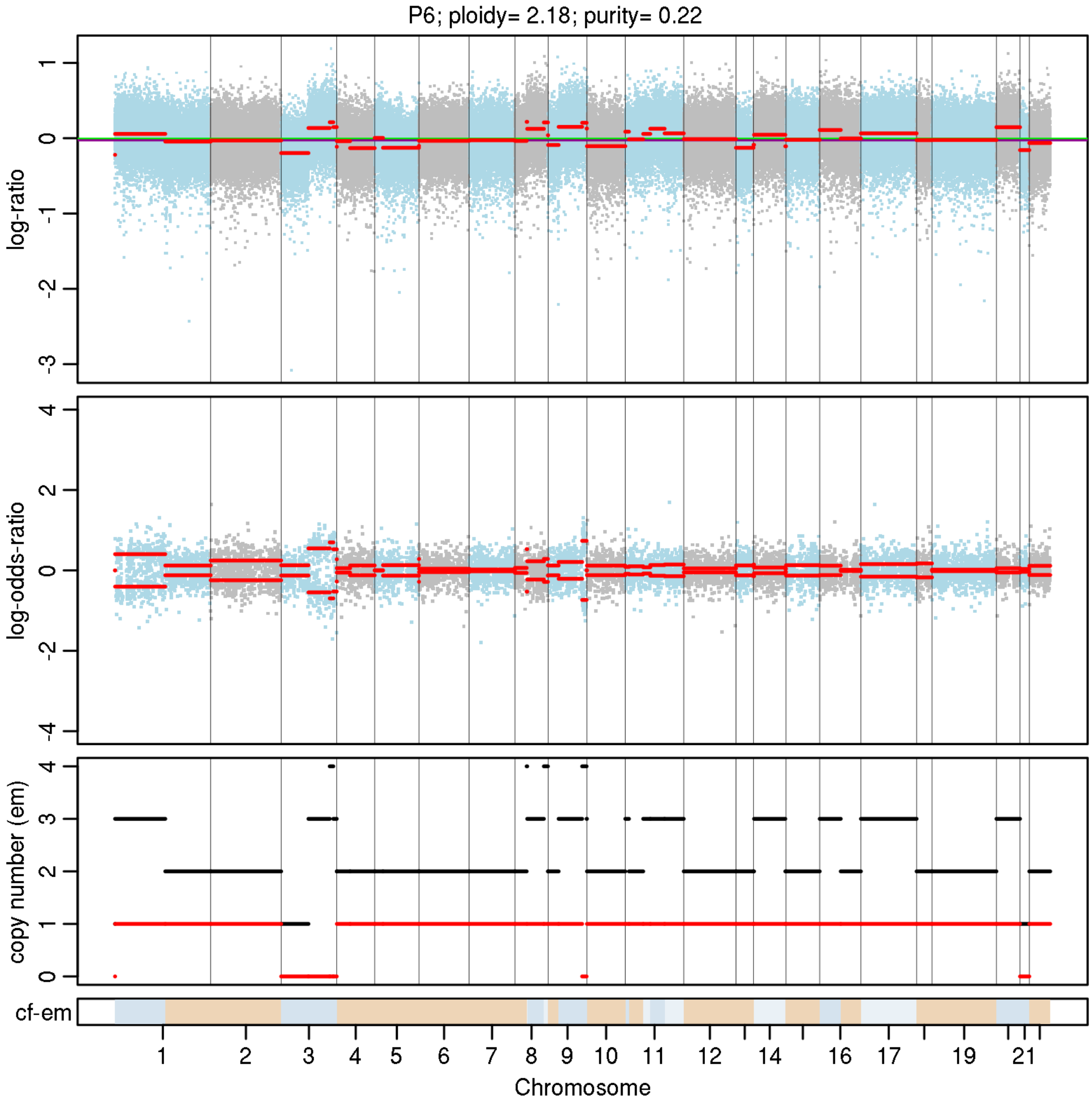

**Supplementary Figure 14. ASCN estimation from patient 6 WES dataset using FACETS.**
The top panel displays logR with chromosomes alternating in blue and gray. The green line indicates the median logR in the sample. The purple line indicates the logR of the diploid state. The second panel displays logOR. Segment means are plotted in red lines. The third panel plots the total (black) and lesser/minor (red) copy numbers for each segment. The bottom bar shows the associated cellular fraction (cf). Darker blue shade indicates higher cf. Beige indicates a normal segment (total=2, minor=1). The entire plot has been generated from the FACETS pipeline implemented through cnv_facets. Refer to the original article [11] for additional details.

### 5. Supplementary Tables

**Supplementary Table 1. ASCNs of CNV-harboring regions estimated by SPICE for colon cancer Slide-seqV2 dataset.**

| Arm | Subclone 1 | Subclone 2 |
|---|---|---|
| 1q | 1,1 | 1,2 |
| 7p | 1,1 | 1,2 |
| 8p | 1,1 | 0,1 |
| 8q | 1,2 | 1,2 |
| 9p | 1,1 | 1,2 |
| 12p | 1,1 | 0,1 |
| 15q | 0,1 | 0,1 |
| 17p | 0,1 | 0,1 |
| 18p | 0,0 | 0,0 |
| 18q | 0,1 | 0,1 |
| 20p | 1,2 | 1,1 |
| 20q | 1,2 | 1,2 |

**Supplementary Table 2. ASCNs of CNV-harboring regions estimated by SPICE for patient 6 Visium dataset.**

| Arm | Subclone 1 | Subclone 2 |
|---|---|---|
| 1p | 1,1 | 1,2 |
| 3q | 1,1 | 1,2 |
| 9q | 1,1 | 1,2 |
| 13q | 1,1 | 0,1 |
| 17p | 1,1 | 1,2 |
| 20p | 1,1 | 1,2 |
| 20q | 1,1 | 1,2 |

**Supplementary Table 3. Summary of CNV-harboring regions estimated by FACETS for patient 6 WES dataset.**

Chromosomes serving as true CNVs were considered for this table. Segments, where only the total copy numbers were inferred, and segments spanning less than 1MB were excluded. TCN: total copy number, LCN: lesser/ minor copy number, Tot.SNP: Number of SNPs in the segment, Het: Number of SNPs that are deemed heterozygous.

| Chr | Start | End | Length | TCN | LCN | Tot.SNP | Het | Arm |
|---|---|---|---|---|---|---|---|---|
| 1 | 786486 | 119915381 | 119128896 | 3 | 1 | 12048 | 517 | 1p |
| 3 | 197709 | 89479453 | 89281745 | 1 | 0 | 6549 | 268 | 3p |
| 3 | 93874276 | 178825759 | 84951484 | 3 | 0 | 5115 | 257 | 3q |
| 3 | 178828239 | 186804597 | 7976359 | 4 | 0 | 850 | 37 | 3q |
| 3 | 186853104 | 198157341 | 11304238 | 3 | 0 | 830 | 39 | 3q |
| 8 | 48060755 | 143158946 | 95098192 | 3 | 1 | 3998 | 221 | 8q |
| 8 | 143213309 | 144850734 | 1637426 | 4 | 1 | 969 | 41 | 8q |
| 9 | 68780972 | 134048439 | 65267468 | 3 | 1 | 5644 | 291 | 9q |
| 9 | 134050533 | 137616038 | 3565506 | 4 | 0 | 1067 | 81 | 9q |
| 11 | 45904804 | 61280947 | 15376144 | 3 | 1 | 1699 | 95 | 11p-11q |
| 11 | 61281109 | 74493412 | 13212304 | 3 | 1 | 3438 | 136 | 11q |
| 11 | 74493683 | 134986747 | 60493065 | 3 | 1 | 4599 | 244 | 11q |
| 14 | 20292119 | 105644629 | 85352511 | 3 | 1 | 7500 | 415 | 14q |
| 17 | 156325 | 83135731 | 82979407 | 3 | 1 | 13370 | 729 | 17p-17q |
| 20 | 87746 | 64273454 | 64185709 | 3 | 1 | 5665 | 319 | 20p-20q |

9. Loh P-R, Danecek P, Palamara PF, Fuchsberger C, A Reshef Y, K Finucane H, et al. Reference-based phasing using the Haplotype

Reference Consortium panel. Nat Genet. 2016;48:1443–8. https://doi.org/10.1038/ng.3679
